# Faster rates of molecular sequence evolution in reproduction-related genes and in species with hypodermic sperm morphologies

**DOI:** 10.1101/2021.08.16.456242

**Authors:** R. Axel W. Wiberg, Jeremias N. Brand, Lukas Schärer

**Affiliations:** University of Basel, Department of Environmental Sciences, Zoological Institute, Vesalgasse 1, 4051 Basel, Switzerland; Max Planck Institute for Biophysical Chemistry, Department of Tissue Dynamics and Regeneration, Am Fassberg 11, 37077 Göttingen, Germany

**Keywords:** Sexual conflict, sexual selection, comparative genomics, transcriptomics, *dN/dS*

## Abstract

Sexual selection drives the evolution of many striking behaviours and morphologies, and should leave signatures of selection at loci underlying these phenotypes. However, while loci thought to be under sexual selection often evolve rapidly, few studies have contrasted rates of molecular sequence evolution at such loci across lineages with different sexual selection contexts. Furthermore, work has focused on separate sexed animals, neglecting alternative sexual systems. We investigate rates of molecular sequence evolution in hermaphroditic flatworms of the genus *Macrostomum*. Specifically, we compare species that exhibit contrasting sperm morphologies, strongly associated with multiple convergent shifts in the mating strategy, reflecting different sexual selection contexts. Species donating and receiving sperm in every mating have sperm with bristles, likely to prevent sperm removal. Meanwhile, species that hypodermically inject sperm lack bristles, potentially as an adaptation to the environment experienced by hypodermic sperm. Combining functional annotations from the model, *M. lignano*, with transcriptomes from 97 congeners, we find genus-wide faster sequence evolution in reproduction-related *versus* ubiquitously-expressed genes, consistent with stronger sexual selection on the former. Additionally, species with hypodermic sperm morphologies had elevated molecular sequence evolution, regardless of a gene’s functional annotation. These genome-wide patterns suggest reduced selection efficiency following shifts to hypodermic mating, possibly due to higher selfing rates in these species. Moreover, we find little evidence for convergent amino acid changes across species. Our work not only shows that reproduction-related genes evolve rapidly also in hermaphroditic animals, but also that well-replicated contrasts of different sexual selection contexts can reveal underappreciated genome-wide effects.

## Introduction

Sexual selection and the various associated processes (e.g. sperm competition, mate choice, and sexual conflict) are potent forces of diversifying selection, driving the evolution of behavioural, morphological, and biochemical traits in many organisms (Andersson 1994; Arnqvist and Rowe 2005). These forces can also be important drivers of population divergence and speciation, due to ensuing rapid evolution and coevolution resulting from intense competition over reproductive resources (Lande 1981; Gavrilets 2000; Hayashi et al. 2007). Understanding the genomic manifestations and consequences of sexual selection has been a long-standing goal in evolutionary biology. For example, sex chromosome evolution is expected to be driven, to a large extent, by sexual conflict (e.g. Frank and Patten 2019), and sex-biased gene expression is thought to result from sexual selection on the underlying sexually dimorphic traits (Grath and Parsch 2016). At the same time, sex-biased genes tend to show rapid rates of molecular sequence evolution (Ellegren and Parsch 2007). But the study of the genomics of sexual selection remains in its infancy, partly because it is difficult to distinguish between the processes of sexual selection, natural selection, and genetic drift based on patterns in sequence data alone (Wilkinson et al. 2015). To make progress, more detailed functional information on the loci in question is needed, as are replicated contrasts between species and populations that differ in how sexual selection operates (Wilkinson et al. 2015).

### Rapid molecular sequence evolution of reproduction-related genes

Reproduction-related genes are defined in several different ways. For example, Swanson and Vacquier (2002) define “reproductive proteins” as “…those that act after copulation and that mediate gamete usage, storage, signal transduction and fertilization”. Meanwhile, Dapper and Wade (2020) define a “reproductive gene” as “…any protein coding-gene with a direct role in any reproductive process.” A broad definition can thus be given as genes linked to some aspect of reproduction, similar to that adopted by Dapper and Wade (2020). This definition includes those involved in conserved reproductive functions, such as meiosis and gamete production (which may or may not be subject to sexual selection), but also any genes that underlie variation in sexually-selected traits (e.g. ornaments, preferences or biases, and persistence/resistance traits). A common way to identify at least a subset of such genes is to focus on genes expressed in various reproductive tissues (e.g. testes, ovaries, accessory glands, and reproductive tracts). In principle, all reproduction-related genes, insofar as they are under sexual selection, could show evidence of positive selection (Swanson and Vacquier 2002; Dapper and Wade 2020).

As expected from models of sexual selection, large-scale studies often show elevated rates of molecular sequence evolution in reproduction-related genes when compared to the rest of the genome (for reviews see for example Swanson and Vacquier 2002; Vacquier and Swanson, 2011; Wilburn and Swanson, 2016). This is often assessed by analysis of the rates of molecular sequence evolution, e.g. by computing the ratio of non-synonymous (*dN*) and synonymous (*dS*) substitution rates in protein-coding DNA sequences (Yang and Bielawski 2000; see below for more details). Rapid evolution has been observed for a range of such genes in several taxa. For example, across mammals, genes governing sperm-egg interactions show strong evidence of positive selection at a few amino acid sites (Swanson et al. 2001). Additionally, in mice, ejaculate proteins and proteins isolated from the seminal vesicle have high rates of molecular sequence evolution that are indicative of positive selection (Ramm et al. 2008; Dean et al. 2009). Meanwhile, also in mice, proteins isolated from other reproduction-related tissues, such as the prostate, show lower rates compared to the genome-wide background, suggesting sequence conservation for some subsets of reproduction-related genes (Dean et al. 2009). High rates of molecular sequence evolution have also been found for proteins involved in sperm-egg interactions in several marine invertebrates (Clark et al. 2009; Aagaard et al. 2013; Vacquier and Swanson 2011). In comparative studies of *Drosophila*, some subsets of reproduction-related genes (namely genes expressed in the testes and accessory glands), are among the fastest evolving genes in the genome, both in terms of sequence divergence and gene gain/loss (Haerty et al. 2007; Findlay et al. 2008; Patlar et al., 2020).

To date, however, few studies have contrasted molecular sequence evolution in closely related lineages that differ in the way in which sexual selection manifests itself. Even fewer studies have taken a wide comparative genomics approach, be it by investigating many genes or a large sample of species. Instead, studies are generally limited either by their focus on only a few genes or by sampling only a few species (e.g. Dorus et al. 2004; Hurle et al. 2007; Ramm et al. 2008; Vicens et al. 2014; Harrison et al. 2015). Furthermore, almost no work has been done to generalise findings to different sexual systems. For example, very few data are available from simultaneous hermaphrodites, organisms where each individual is both male and female at the same time, especially in animals. Some recent studies of hermaphrodites exist from plant systems, but these generally also focus on only a few species (Szövényi et al. 2013; Arunkumar et al. 2013; Gossmann et al. 2013; 2016; Harrison et al. 2019; Moyle et al. 2021), and comparisons between plants and animals are challenging for various reasons. For example, the effect of haploid selection might be important in plants but negligible in animals (Joseph and Kirkpatrick 2004, but see Alavioon et al. 2017).

In addition to the limitations above, there is often a difficulty in drawing inferences from observed patterns to the underlying processes. Signatures of rapid molecular sequence evolution are typically taken as evidence for strong sexual selection acting on reproduction-related genes. However, relaxed selection on coding sequences, either due to reduced intensity or reduced efficiency of selection, can also result in accelerated molecular sequence evolution (Wertheim et al. 2015; Dapper and Wade 2016; 2020). Indeed, there are several reasons why selection may in fact be less efficient on reproduction-related genes than other genomic loci (Dapper and Wade 2016; 2020). For example, reproduction-related genes are often sex-biased, or even sex-limited, in their expression (Dapper and Wade 2016; 2020). In the case of sex-limited expression, such genes are “visible” to selection only in one sex and, all else being equal, the selection they experience is therefore expected to be only half that of genes expressed equally in both sexes (Dapper and Wade 2016; 2020). It may therefore not be appropriate to compare reproduction-related genes to a genomic background set of genes, and more nuanced null models should be considered (Dapper and Wade 2016; 2020). However, such null models have, to date, only been developed for species with separate sexes, and theory about how they might apply to, for example, simultaneous hermaphrodites is currently lacking. Every simultaneous hermaphrodite is both male and female at the same time. It would therefore seem that a gene cannot, on average, be sex-biased or sex-limited in expression in a sense that would produce the kind of consistent differences in the efficiency of selection highlighted by Dapper and Wade (2016; 2020). As a result, reproduction-related genes in simultaneous hermaphrodites may not experience lower efficiency of selection compared to other genes. Therefore, in hermaphrodites any differences in the rates of molecular sequence evolution between reproduction-related genes and other genes may be more easily attributable to differences in the actual intensity of selection. This potentially makes simultaneous hermaphrodites, including the genus *Macrostomum* we introduce below, powerful systems to study the evolution of reproduction-related genes and the effects of sexual selection on the genome. It is clear that more work is sorely needed to probe the consequences of different sexual selection contexts and sexual systems on the genes underlying sexually-selected traits and on the genome as a whole. At the same time, it is important to apply an analysis framework that allows testing of competing hypotheses to explain patterns of molecular sequence evolution.

Estimating the effects of evolutionary forces on molecular sequences is a challenging task and relying on a single approach is problematic. As noted above, the effect of selection on molecular sequence evolution is often assessed by computing the ratio of non-synonymous (*dN*) to synonymous (*dS*) substitution rates in protein-coding DNA sequences (*dN/dS* = ω; Yang and Bielawski 2000). If synonymous substitutions, at the coding sequence level, are assumed to be neutral, then a value of ω = 1 suggests that a gene is evolving neutrally, since both types of substitution occur at the same rate. If ω < 1 or > 1 a gene is inferred to be evolving under purifying or positive selection, respectively, since non-synonymous substitutions occur less or more frequently than expected. Maximum-likelihood methods for the computation of ω using codon-substitution models are readily available. “Branch models” estimate a gene-wide ω to contrast average ω values for genes across different lineages in a phylogeny (Yang and Bielawski 2000). While flexible and useful, these models have some limitations. For instance, even if positive selection is acting on a gene, this is unlikely to raise ω > 1 for the entire coding sequence. While changes may be favoured in a specific binding region, the overall protein backbone may still be conserved. Thus, most genes typically show values of ω < 1. “Branch-site models”, in turn, permit more complex scenarios where most sites are under purifying selection while allowing a few sites to be under positive selection (Yang and Bielawski 2000). While there is ongoing work on how robust branch-site tests are to violations of model assumptions (e.g. Fletcher and Yang 2010; Yang and dos Reis 2011; dos Reis and Yang 2013; Gharib and Robinson-Rechavi 2013; Venkat et al. 2018), the consensus suggests that it is a fairly robust approach. Additionally, because a shift from purifying to relaxed selection can result in similar values of ω to those expected from positive selection, especially in branch models, it is prudent to also explicitly test for evidence of relaxed selection. Specifically, branch-site models can be employed to test if the differences in the distributions of ω values across sites between a set of test and reference branches in a phylogeny are consistent with expectations from a relaxation of selection in test branches (Wertheim et al. 2015). In summary, using multiple methods with different assumptions and approaches allows for a more complete picture of the role of selection in shaping patterns of sequence divergence.

### Reproductive biology of the genus Macrostomum

Free-living flatworms of the species-rich genus *Macrostomum* are members of Macrostomorpha, a large group of Platyhelminthes, representatives of which vary tremendously in reproductive traits (Janssen et al. 2015). *Macrostomum* are simultaneous hermaphrodites, producing sperm and eggs at the same time, and several species from this genus, particularly the main model species *M. lignano* (Ladurner et al. 2005; Wudarski et al. 2020), are used to study mating behaviour, sexual selection, and sexual conflict (e.g. Schärer et al. 2005; Ramm et al. 2015; Marie-Orleach et al. 2017; Giannakara et al. 2016; Giannakara and Ramm 2017; Patlar et al. 2020; Singh et al. 2020a, 2020b). In hermaphrodites, sexual conflict is expected to occur not just over the mating and re-mating rates, but also over the sex roles that the partners assume during copulation, with the male (or “sperm donor”) role generally thought to be preferred over the female (or “sperm recipient”) role (Charnov 1979; Michiels 1998; Schärer et al. 2015). These conflicts have likely been important in driving the evolution of the extraordinary diversity in reproductive morphologies and behaviours seen in this genus (Schärer et al. 2011).

Many *Macrostomum* species employ a strategy of reciprocal copulation (figure 1A), where both partners donate and receive sperm at the same time, by reciprocally inserting their male intromittent organ, the stylet, into the partner’s sperm receiving organ, the antrum (Schärer et al. 2004; Vizoso et al. 2010). In contrast, other species employ a strategy of hypodermic insemination (figure 1A), where individuals inject sperm into the body of the partner through the epidermis using a needle-like stylet (Schärer et al. 2011). These two mating strategies are associated with characteristic differences in stylet and sperm morphology (Schärer et al. 2011). Specifically, reciprocal copulation is associated with sperm having stiff lateral bristles (figure 1B), which are thought to serve as anchoring mechanisms preventing sperm removal by the recipient (Vizoso et al. 2010; Schärer et al. 2011). Indeed, in several species it is observed that, after copulation, the recipient will place its mouth opening over its own female genital opening in a stereotypical *sucking* behaviour in an apparent attempt to remove sperm and/or other ejaculate components (Schärer et al. 2004; Vizoso et al. 2010; Schärer et al. 2020). Meanwhile, hypodermic insemination is associated with a simpler sperm morphology lacking these bristles (Schärer et al. 2011; figure 1B). Additionally, some species display an intermediate sperm morphology with reduced bristles (Schärer et al. 2011; figure 1B).

**Figure 1.**
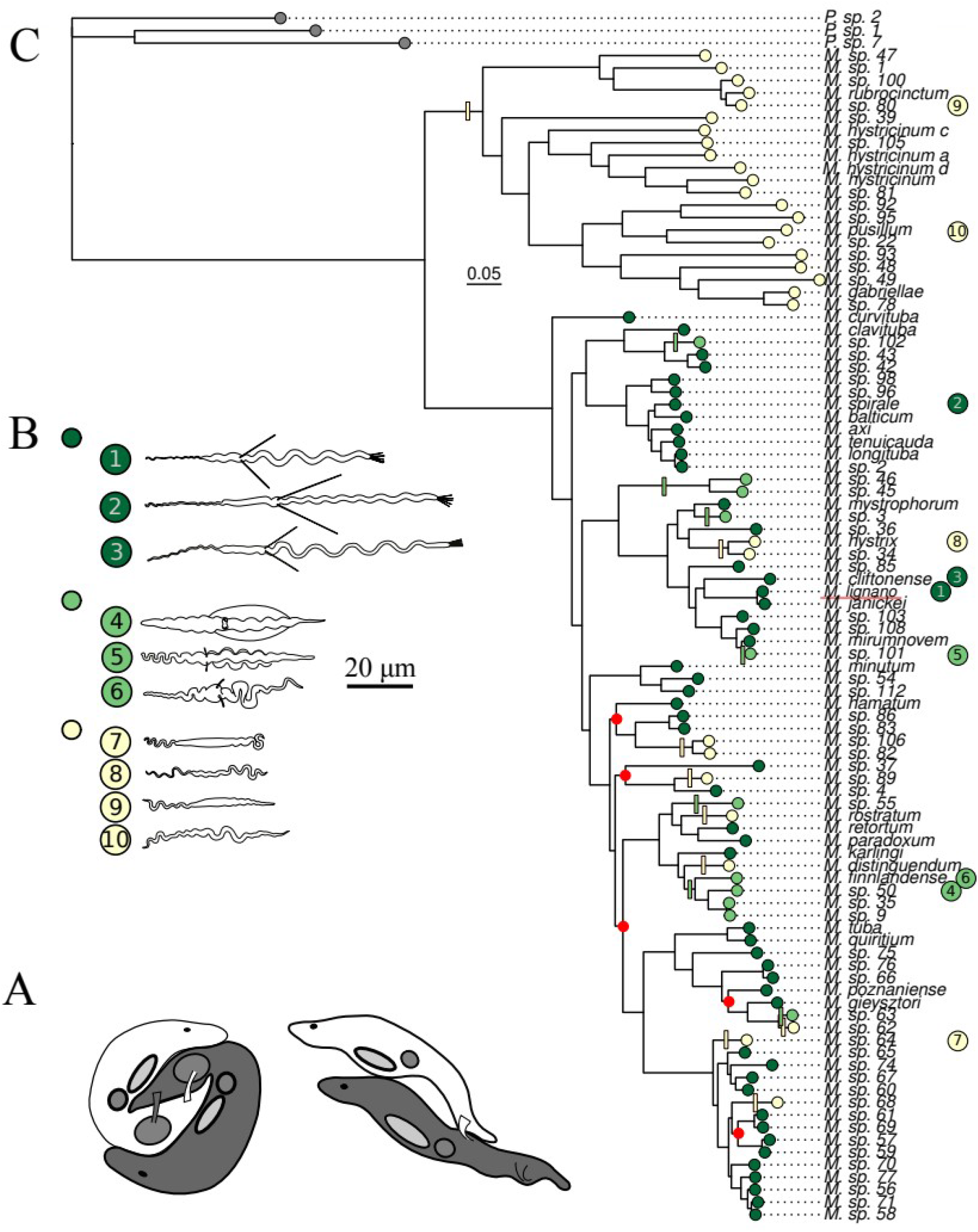
**A** Stylised drawings of *Macrostomum* species with reciprocal copulation (left) and hypodermic insemination (right), respectively, adapted from drawings by Dita Vizoso. **B** Examples of sperm from each bristle state category, from top to bottom: dark green - present, shown are *M. lignano* (1), *M. spirale* (2), and *M. cliftonense* (3); light green - reduced, shown are *M.* sp. 50 (4), *M.* sp. 101 (5), and *M. finnlandense* (6), and yellow - absent, shown are *M.* sp. 64 (7)*, M. hystrix* (8), *M.* sp. 80 (9), and *M. pusillum* (10). **C** Phylogeny of the genus *Macrostomum* based on *de novo* transcriptomes (referred to as H-IQ-TREE in Brand et al,. *in press*), pruned to contain only the 97 species used in this study, as well as three outgroup species of the sister genus *Psammomacrostomum*. The scale bar depicts the number of substitutions per site. Tips are coloured according to the character state of the sperm bristles (see **B**). Coloured bars along branches indicate the phylogenetically independent losses (yellow) and reductions (light green) inferred from ancestral state reconstructions (from Brand et al. *unpublished data*). The outgroup species (grey labels) do not have sperm bristles. Red points indicate nodes with < 90% ultrafast bootstrap support (from Brand et al. *in press*). *M. lignano,* an established model species within this group, is underlined in red.

Recent comparative work on 145 *Macrostomum* species has now greatly expanded the number of known species (Brand et al. *in press*) and investigated the evolution of morphological traits across the genus *Macrostomum* (Brand et al. *unpublished data*). These studies provide evidence for striking convergent evolution, with at least 9 independent losses and 7 independent reductions of sperm bristles across the genus (figure 1C). They also confirm that the contrasting bristle and sperm morphologies are strongly associated with differences in the mating strategies exhibited by the species, by showing that the evolution of hypodermic insemination predicts variation in major axes of morphospace (Brand et al. *unpublished data*). The reproductive morphology can therefore be considered a proxy for the contrasting mating strategies, which represent different outcomes of sexual conflict over sex roles during mating, and probably strongly affect the sexual selection context experienced by sperm and other reproductive traits. For example, a shift to a hypodermic mating strategy might result in a more fair-raffle like sperm competition between the ejaculates of rival individuals that compete within a recipient (Schärer and Janicke 2009; Schärer et al. 2011). A shift in the sperm competition arena, namely from the female reproductive tract to different tissues in the body of the recipient, may also drastically alter the types of sperm traits selected that are favoured. This should also be reflected in patterns of molecular evolution in genes underlying these traits.

### Study overview

Here we combine several sources of information to investigate patterns of molecular sequence evolution of reproduction-related genes in the genus *Macrostomum*. We produce alignments of orthologous genes using *de novo* transcriptome assemblies of ∼100 species of *Macrostomum* available from a recent large-scale phylogenomic study (Brand et al. *in press*; figure 1C). Moreover, annotations for a large number of genes are available from gene expression and functional studies in the model species *M. lignano*. For example, using a positional RNA-Seq approach Arbore et al. (2015) identified transcripts that were primarily expressed in body regions containing specific reproductive tissues, i.e. the testes, the ovaries, and the tail (containing male and female genitalia, as well as prostate gland cells). Further work using *in situ* hybridisation narrowed down transcripts expressed in these regions to those exclusively localised in, for example, the testis, the ovary, the female reproductive tract (the antrum), or the prostate gland cells, thus largely validating these positional annotations (Arbore et al. 2015; Lengerer et al. 2018; Weber et al. 2018; Ramm et al. 2019). Evidence has also been presented that these reproduction-related genes evolve rapidly (Brand et al. 2020), but a crucial limitation of that study is that it investigated only four species and thus had little power to compare lineages with different sexual selection contexts.

In this study, we take advantage of a greatly expanded dataset to investigate the evolution of reproduction-related genes throughout the genus *Macrostomum*. We fit different codon models of molecular sequence evolution introduced above (see also Materials and Methods) to alignments of genes expressed in different body regions that contain important reproductive tissues in *M. lignano* (reproduction-related genes). Our study tests the hypotheses that molecular divergence in these genes are due to positive or relaxed selection. Specifically, we compare reproduction-related genes to genes that are likely ubiquitously expressed throughout the worm. Moreover, we contrast patterns of molecular sequence evolution between species that differ in whether their sperm morphology includes sperm bristles or not, in order to assess the effect of a changing sexual selection context on the evolution of reproduction-related genes.

## Results

### Branch models

We identified a total of 1,149 orthogroup alignments (OGs) that passed all successive steps of the bioinformatic pipeline (see Materials and Methods; figure S1) and thus could be used in analyses of molecular sequence evolution with PAML v. 4.9 (table S1; Yang 2007). Based on RNA-Seq data from the model organism *M*. *lignano* (Arbore et al. 2015), we annotated 892 OGs as likely reproduction-related (yielding 643, 135, and 114 testis-, ovary-, and tail-region OGs, respectively), and 257 as likely showing ubiquitous expression (ubiquitously-expressed OGs). To identify broad differences in the patterns of molecular sequence evolution among these annotation groups, we first fit a codon model of molecular sequence evolution that assumes a single common ω value for all lineages. On the assumption that OGs can be treated as statistically independent, we found a significant effect of annotation group on average ω (Kruskal-Wallis rank-sum test, χ^2^ = 34.93, d.f. = 3, p < 0.001; figure 2A). In *post-hoc* pairwise comparisons we found that this was driven by ubiquitously-expressed OGs having lower average ω, while the other annotation groups showed similar values (figure 2A). Furthermore, we found a negative correlation between the expression level of the *M. lignano* gene in each OG (as measured from the whole body) and the estimated ω value for that OG (Spearman rank correlation, rho = -0.72, p = 0.015). Thus, more highly expressed OGs show slower rates of molecular sequence evolution.

**Figure 2.**
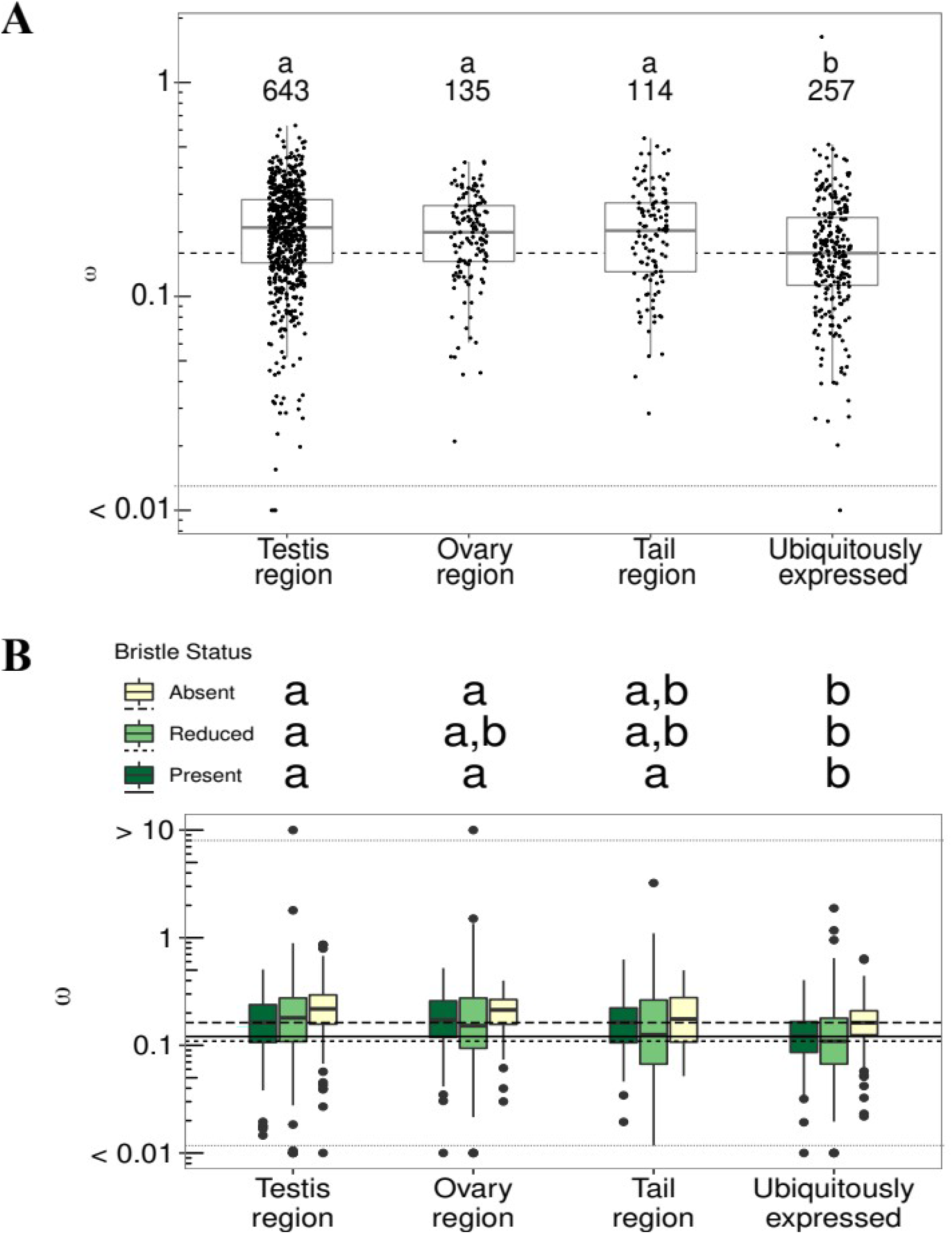
**A** *dN/dS* (ω) values, from a model assuming a single ω value for the whole tree, for orthogroups (OGs) assigned to different (functional) annotation groups, as well as a set of OGs annotated as ubiquitously expressed (out of 1,000 such ubiquitously-expressed OGs initially selected at random). The horizontal dashed line gives the median ω value for the ubiquitously expressed OGs. Sample sizes for each annotation group are given as inset text. **B** *dN/dS* (ω) values across OGs with in different annotation groups for species with different sperm morphologies (bristle state). The solid, short-dashed, and long-dashed horizontal lines give the median ω values across ubiquitously-expressed OGs for species with different sperm morphologies (see inset legend). For both **A** and **B** the letters show nonparametric all-pairs post-hoc tests (groups with different letters are significantly different at an adjusted p-value < 0.05). Note that for ease of plotting, some OGs with ω estimates > 10 or < 0.01 (N = 15) have been fixed to ω values of 10 or 0.01, respectively, and separated from the remaining points with stippled horizontal lines.

We next asked if ω differed across the sperm bristle states, both between and within annotation groups. Specifically, for 828 OGs that contained representative species of all three bristle states, we compared the above single ω models to models with three separate ω values, one for each bristle state. Overall, we found that 514 (∼62%) OGs were better explained by the model with three states (table S1), thus clearly documenting variation in ω across sperm bristle states. Ideally, we would test the two factors “bristle state” and “annotation group” simultaneously, but given the distributional properties of the ω values (i.e. skewed distributions and unequal variances between the bristle states), we in the main text report only separate non-parametric statistics for the differences (i) between annotation groups within each bristle state and (ii) across bristle states within each annotation group (but see also Supplementary Text T1 in the Supplementary Materials for parametric statistics on log_10_() transformed ω values, including an interaction term, which give qualitatively similar results).

For species with bristles present, we found significant differences in ω between annotation groups (Kruskal-Wallis rank-sum test, χ^2^ = 25.2, d.f. = 3, p < 0.001), in that testis-, ovary-, and tail-region OGs showed higher average ω values than ubiquitously-expressed OGs (figure 2B), and similar patterns were observed for species with reduced bristles (χ^2^ = 24.9, d.f. = 3, p < 0.001) or absent bristles (χ^2^ = 32.3, d.f. = 3, p < 0.001). In summary, we found that the pattern of higher ω values for reproduction-related OGs, compared to the ubiquitously-expressed OGs (figure 2A), was recapitulated within the lineages with different bristle states (figure 2B).

Because estimates of ω are arguably more informative when more evolutionary lineages are included, we further checked if these results were robust when we only considered OGs containing sequences from species covering ≥3 phylogenetically independent losses of bristles. Here the only qualitative change was that there was no longer a significant difference between the tail-region OGs and the ubiquitously-expressed OGs for any bristle state (figure S2). Removing OGs with few independent origins greatly reduced the sample sizes, also for the OGs with the already sparse tail-region annotations (from N = 114 to N = 48). Identifying orthologs for tail-region genes might be particularly difficult due to their rapid evolution. Brand et al. (2020) also found that testis-, ovary-, and tail-region OGs were under-represented among OGs that contained representative sequences from all four species in their study, indicating that homologs were more difficult to detect for these genes. The above differences among annotation groups were also qualitatively the same if we removed low-expressed OGs (figure S3) or if we removed some OGs containing residual cDNA synthesis primer sequences (see Supplementary Text T2 in the Supplementary Materials).

Next, we compared ω across the different bristle states within annotation groups. Although this effect is already evident in figure 2B, for visualisation it is helpful to split the data according to different patterns observed among the bristle states. When we split the data by whether ω decreases (–) or increases (+) as one moves from species with bristles to those with absent bristles (figure 3), we observed that ω values are more often higher in species with absent bristles (i.e. the right panel contains more OGs). Among the testis- and ovary-region OGs we found a significant overall difference among the three bristle states (Kruskal-Wallis rank-sum tests, testis-region, χ^2^ = 42.2, d.f. = 2, p < 0.001; ovary-region, χ^2^ = 7.3, d.f. = 2, p =0.03), while tail-region OGs showed a non-significant pattern in the same direction (χ^2^ = 3.5, d.f. = 2, p = 0.18), and also the highest percentage of OGs with the opposite pattern (32.6%, figure 3, left panel).

**Figure 3.**
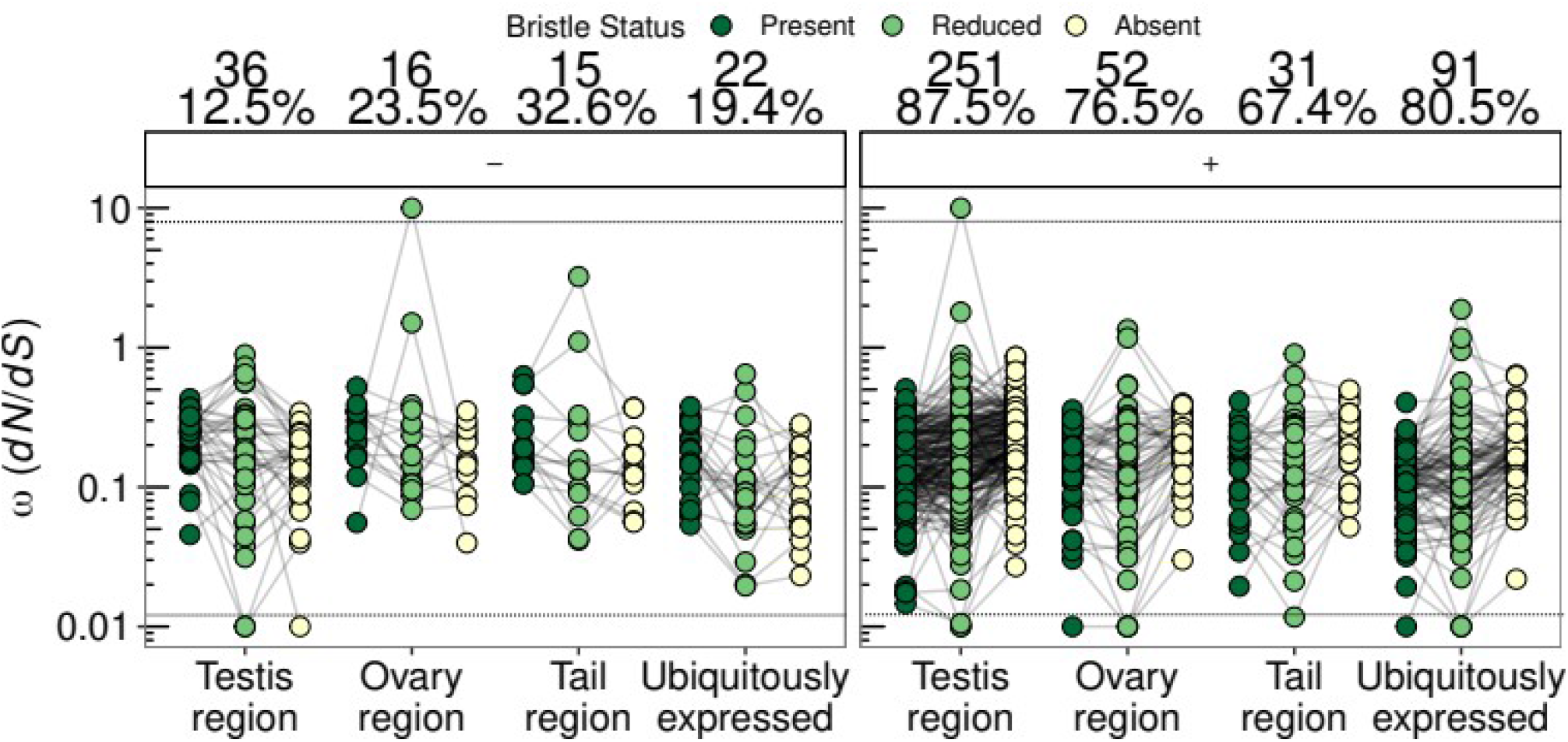
*dN/dS* (ω) values for orthogroups (OGs) that were better explained by a model assuming different ω values for species with the three contrasting sperm morphologies (bristle states). Values for the same OG in species with different bristle states are connected by a line. Panels show the data split into different patterns, namely with ω decreasing (–, left panel) or increasing (+, right panel) as one moves from species with bristles to those lacking bristles. The text above each panel gives the number and percentage of OGs across annotation groups and patterns of ω. Note that for ease of plotting, some OGs with ω estimates > 10 or < 0.01 have been fixed to ω values of 10 or 0.01, respectively, and separated from the remaining points with stippled horizontal lines.

Intriguingly, we found that this overall pattern was true also for ubiquitously-expressed OGs (χ^2^ = 19.1, d.f. = 2, p < 0.001), indicating that the higher ω for species with absent sperm bristles appears to be a genome-wide characteristic, rather than one that is specific for reproduction-related OGs. Pooling data across all annotation groups, we found an even stronger effect of bristle state (χ^2^ = 59.1, d.f. = 2, p < 0.001). Across all annotation groups, the majority (67.4-87.5%) of OGs had higher ω values in species with absent bristles (figure 3). Moreover, testis-region OGs had the highest percentage of OGs with higher ω among species with absent bristles (87.5%; figure 3, right panel). However, there was no significant overall deviation from the proportions expected based on the distribution of all tested OGs in these annotation groups (χ^2^-test, χ^2^ = 4.6, d.f. = 3, p = 0.2).

We also observed that the variance in the estimates of ω across OGs was highest for the lineages with reduced bristles (figure 3). This was even more apparent when we visualized the data split into four different patterns of the ω values across the three bristle states (see the four panels labelled –/–, –/+, +/–, and +/+ in figure S4). The patterns –/– and +/+ are, respectively, interpretable as ω always decreasing (–) or increasing (+) as one moves from species with bristles present, *via* those with reduced bristles, to those with absent bristles. The other two patterns (–/+ and +/–) reveal that species with reduced bristles sometimes do not show intermediate ω values. Deviations from the –/– and +/+ pattern could occur if species with reduced bristles do not represent a straightforward transitional state between present and absent bristles. In this case, the peculiarities of mating strategies within this reduced bristle state may impose different selection pressures on reproduction-related genes for some species, leading to increased variation within this state. Alternatively, a higher variance in estimates of ω for the reduced bristles category could arise due to the generally lower number of representatives for this bristle state in our data, both in terms of their absolute numbers (figure S5A), and in terms of the represented number of phylogenetically independent losses compared to reductions of the sperm bristles (figure S5B and C), thus potentially making estimation less reliable.

We also confirmed that these results were robust to various potentially confounding factors. Visual inspection of the data suggested that the OGs that were better explained by a model with three ω values compared to a single ω value did not show strong biases in terms of average pairwise *dS* (figure S6A) or overall tree length (figure S6B). Thus, we concluded that *dS* saturation was not driving these results. Moreover, systematic differences in codon usage bias between lineages with contrasting sperm morphologies could also give spurious signals of positive selection. However, OGs did not show evidently different levels of codon usage bias (figure S6C). Nor were OG alignment lengths different (figure S6D), suggesting that a simple difference of power stemming from longer sequences (and the correspondingly greater opportunity for mutations to occur and fix) was not driving these results. Finally, across the annotation groups, 42.1-54.1% of OGs used in this analysis contained representatives covering ≥3 independent losses of sperm bristles (figure S5B) and 24.6-45.2% of OGs contained representatives covering ≥3 independent reductions of sperm bristles (figure S5C). Thus, in many cases our estimates were derived from several independent transitions, giving us a greater confidence in them.

### Branch-site models

While branch models give an informative measure of the relative rates of molecular sequence evolution, they are not ideal for detecting strong positive selection acting on only a few sites within a landscape of overall purifying selection (Zhang et al. 2005). We therefore also applied branch-site models to a subset of the OGs (see also Materials and Methods for details). Briefly, we subset the data to perform contrasts for species with bristles present compared to species with absent bristles (present-absent) and for species with bristles present compared to species with reduced bristles (present-reduced). In both cases we considered species with bristles present as the background branch. We were mainly interested in these two contrasts because present-absent represents a comparison between the hypothesised start and endpoints of the transition in morphology (and in mating strategy), while the present-reduced contrast could represent early stages of transition (Brand et al. *unpublished data*). We fit the branch-site model A (Zhang et al. 2005), which allows for positive selection (ω > 1) among branches labelled as foreground, and compare this to the corresponding null model (A_null_) that is otherwise identical, but does not allow for positive selection. Thus, for a given OG, a significantly better fit of model A and a ω estimate > 1 can be taken as evidence of positive selection acting on some subset of sites within the gene (see the Materials and Methods for more details).

We found that relatively few OGs showed strong evidence for having sites evolving under positive selection in either of these contrasts (table 1, table S2), compared to the proportion of OGs with evidence for different ω values depending on the bristle state (see *Branch models* above). In the present-absent and present-reduced contrast, we found that only a total of 64 (6.8%) and 73 (7.6%) OGs showed evidence of positive selection (i.e. LRT p-values < 0.05), respectively. However, in neither contrast was there a clear difference in the ω_2_ parameter (see Materials and Methods) across the annotation groups (Kruskal-Wallis rank-sum tests, present-absent, χ^2^ = 6.6, d.f. = 3, p = 0.09; figure S7A; present-reduced, χ^2^ = 1.0, d.f. = 3, p = 0.81; figure S7B). Furthermore, in neither contrast did we find a clear deviation of the distribution of significant OGs from that expected based on the distribution of all OGs across annotation groups (χ^2^-tests, present-absent, χ^2^ = 3.3, d.f. = 3, p = 0.35; present-reduced, χ^2^-test, χ^2^ = 3.3, d.f. = 3, p =0.34; table 1, table S2). These results suggest that in both contrasts all annotation groups are equally likely to contain genes inferred to contain sites under positive selection and there is no clear evidence for more intense selection in any one annotation group. Finally, a large proportion of the alignments with evidence for positive selection represent alignments that contain only a single loss or reduction of bristles (present-absent, 64%; present-reduced, 47.9%), although in both contrasts some alignments contained up to 7 and 4 independent shifts, respectively (table S2).

**Table 1.** Analyses of molecular sequence evolution using branch-site models. Total number of orthogroups (OGs) tested (in square brackets), and the number and percentage (in brackets) of OGs in each annotation group for which a branch-site model—allowing for specific sites to be under positive selection in foreground compared to background branches—represents a better fit (LRT p-value < 0.05). Results are shown for both contrasts tested, namely species with bristles present *vs.* species with bristles absent (present-absent), and species with bristles present *vs*. species with reduced bristles (present-reduced).

| Annotation group | present-absent | present-reduced |
| --- | --- | --- |
|  | **Significant (%)**<br>**[total]** | **Significant (%)**<br>**[total]** |
| **Testis region** | 38<br>(7.6)<br>[503] | 41<br>(7.4)<br>[555] |
| **Ovary region** | 6<br>(5.2)<br>[116] | 7<br>(5.8)<br>[121] |
| **Tail region** | 9<br>(9.5)<br>[95] | 11<br>(12.3)<br>[89] |
| **Ubiquitously expressed** | 11<br>(4.7)<br>[231] | 14<br>(7)<br>[201] |
| **Total** | 64<br>(6.8)<br>[945] | 73<br>(7.6)<br>[966] |

Based on posterior probabilities that ω > 1 for each site (Yang et al. 2005; see Materials and Methods for more details), relatively few genes showed evidence for positive selection in branch-site tests. Full details of which genes/sites are inferred to be under positive selection in the present-absent and present-reduced contrasts are given in table S2, and a selection of alignments is highlighted in figure S8. In those alignments with evidence for positive selection it was often the case that only a single foreground branch was driving this result in an otherwise good alignment (e.g. figure S8A). One should, therefore, arguably place more weight on signals from well-resolved alignments, containing multiple losses/reductions of sperm bristles, where amino acid changes have occurred in several foreground species within long stretches of well-aligned sequence (e.g. figure S8B and C), and not necessarily on those with the lowest p-values or those that pass corrections for multiple testing. We partially avoided this by stratifying our analysis approach by the number of independent losses/reductions (see Materials and Methods). In other alignments, probably the majority, some parts of the alignment are fragmented, but the sites inferred to be under selection lie within relatively well-aligned regions (e.g. figure S8D). Finally, in some cases the sites inferred to be under positive selection lay in regions of fragmented sequences from species with either absent or reduced bristles (e.g. figure S8E, F, G). Robust inferences are difficult in the last category of alignments, and the estimated values should be taken with some scepticism. However, although such fragmented regions may, in part, be due to incomplete transcripts in the *de novo* transcriptome assemblies we have used here, it is also possible that they represent biological reality. For example, occasionally the foreground species have a very different sequence structure from other species (e.g. figure S8F). Such cases may represent structural evolution, be it adaptive or neutral, of molecular sequences, which is not easily captured by the codon substitution models we employed here. Identification of sites under selection therefore often requires careful examination of alignments (which we therefore provide in full in Supplementary Archive 1, available at: https://doi.org/10.5281/zenodo.4972188), although this naturally becomes difficult in large-scale studies involving hundreds or thousands of alignments.

### RELAX models

The genome-wide elevated ω values in lineages with absent or reduced bristles implied a strong overall difference in selective regime. It seemed unlikely that this could entirely be attributed to positive selection. Instead, the increase in ω could be due to a more general process such as relaxed selection. We, therefore, used RELAX (Wertheim et al. 2015) to test whether some of the elevated ω values were consistent with relaxed selection in foreground branch species. RELAX compares the distributions of ω values in test *vs.* reference branches, by fitting a model where test branches have a distribution of ω values different from that of reference branches according to an exponential function with a parameter *k*, which is interpreted as a “selection intensity parameter” (Wertheim et al. 2015). A value of *k* > 1 indicates intensified selection, and a value of *k* < 1 indicates relaxed selection. This alternative model is compared to a null model that is otherwise identical, but where *k* is fixed to 1 (see Materials and Methods for details). In the analyses performed here the species with bristles present were always used as the reference branch. For visualization, we also fit the less constrained Partitioned Descriptive Model (PDM) to the data (see Materials and Methods for details).

In the present-absent contrast, a larger proportion of OGs showed evidence for relaxed (k<1) selection (32.1%), than for intensified (*k* > 1) selection (6.8%; table 2). Overall, the proportion of OGs with a significantly different distribution of ω values varied across the annotation groups, deviating significantly from those expected from the distribution of all OGs tested across the annotation groups, in that there were more observed OGs with the testis-region annotation (239) than expected (195; χ^2^-test, χ^2^ = 21.6, d.f. = 3, p < 0.001; table 2). However, when we split the data into OGs with *k* > 1 and *k* < 1, we only found a significant deviation from expected distributions for the latter (*k* > 1, χ^2^ = 2.7, d.f. = 3, p = 0.43; *k* < 1, χ^2^ = 31.2, d.f. = 3, p < 0.001; table 2). We also found that both for OGs with *k* > 1 and *k <* 1 there was no significant difference in the *k* parameter across the annotation groups (Kruskal-Wallis rank-sum tests, *k* > 1, χ^2^ = 0.8, d.f. = 3, p = 0.85; *k <* 1, χ^2^ = 2.4, d.f. = 3, p = 0.49; figure 4A). Thus, we found that relaxed selection was relatively common and that OGs annotated as testis-region specific were more likely to be evolving by relaxed selection in the species with absent bristles than expected. However, we found that there was no clear difference in the degree to which selection is relaxed or intensified across the annotation groups (see figure S9 for examples of how distributions of ω covary with *k* estimates). To better understand the changes in the distributions of ω, we further chose to visualize the OGs with the 5 highest and 5 lowest values of *k,* as well as 20 random OGs for closer examination (figure S10). From figure S10 it is evident that where *k* > 1, most sites were actually under more intense purifying selection in the test branches rather than under increased positive selection.

**Figure 4.**
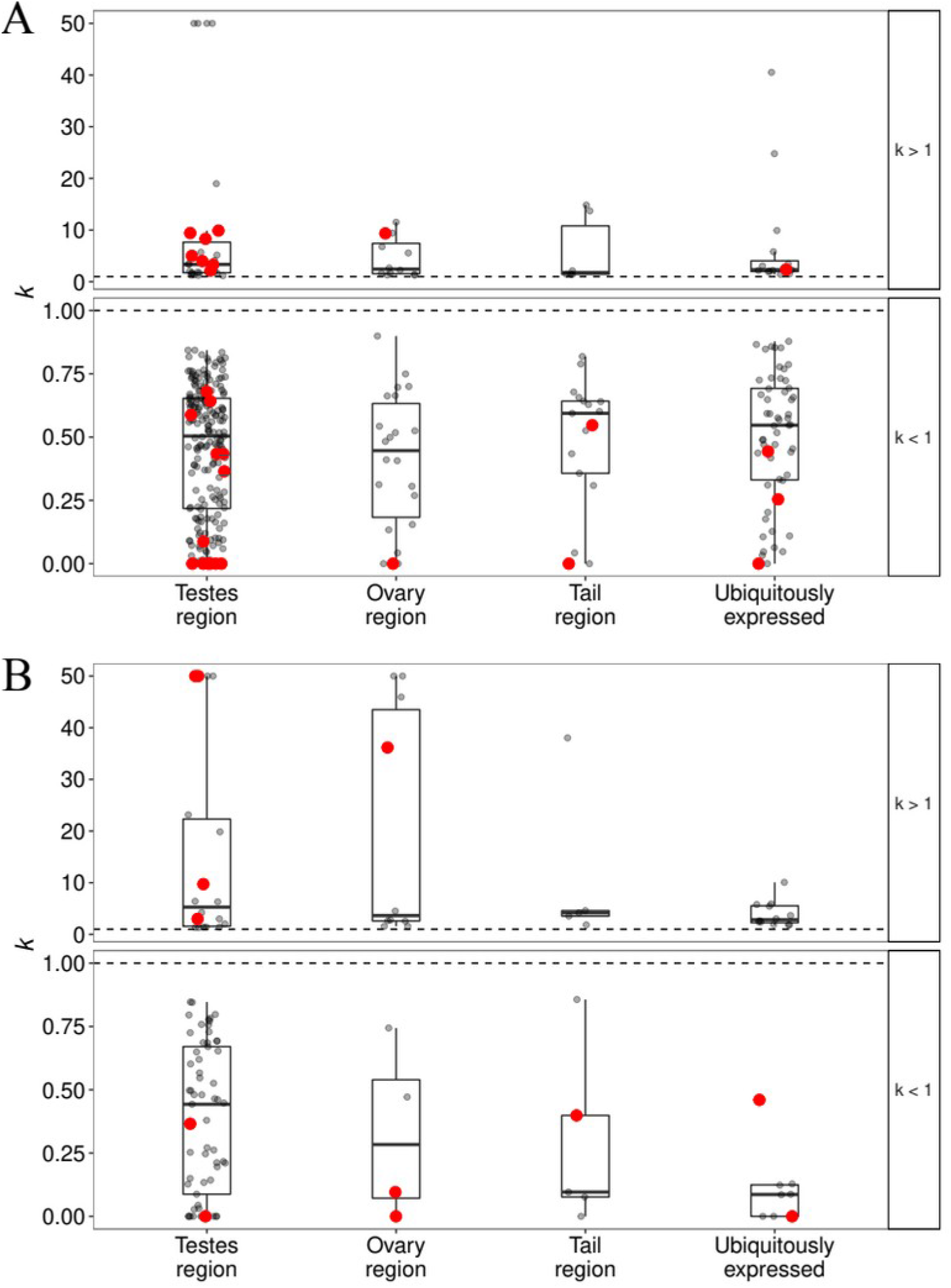
*k* values for orthogroups (OGs) in different (functional) annotation groups from the contrast of species with bristles present and species with absent bristles (**A**), and the contrast of species with bristles present and species with reduced bristles (**B**). In both **A** and **B**, plotted values are restricted to those that show a significant difference in the distribution of ω values between test and reference branches. Additionally, data are split by whether the test showed evidence for intensified (*k* > 1) and relaxed (*k* < 1) selection, respectively. The horizontal black dashed line gives as a reference the *k = 1* line, where the distribution of ω values would be considered the same between test and reference branches. Red points highlight OGs that also showed evidence for positive selection among species with absent bristles in a branch-site models.

**Table 2.**
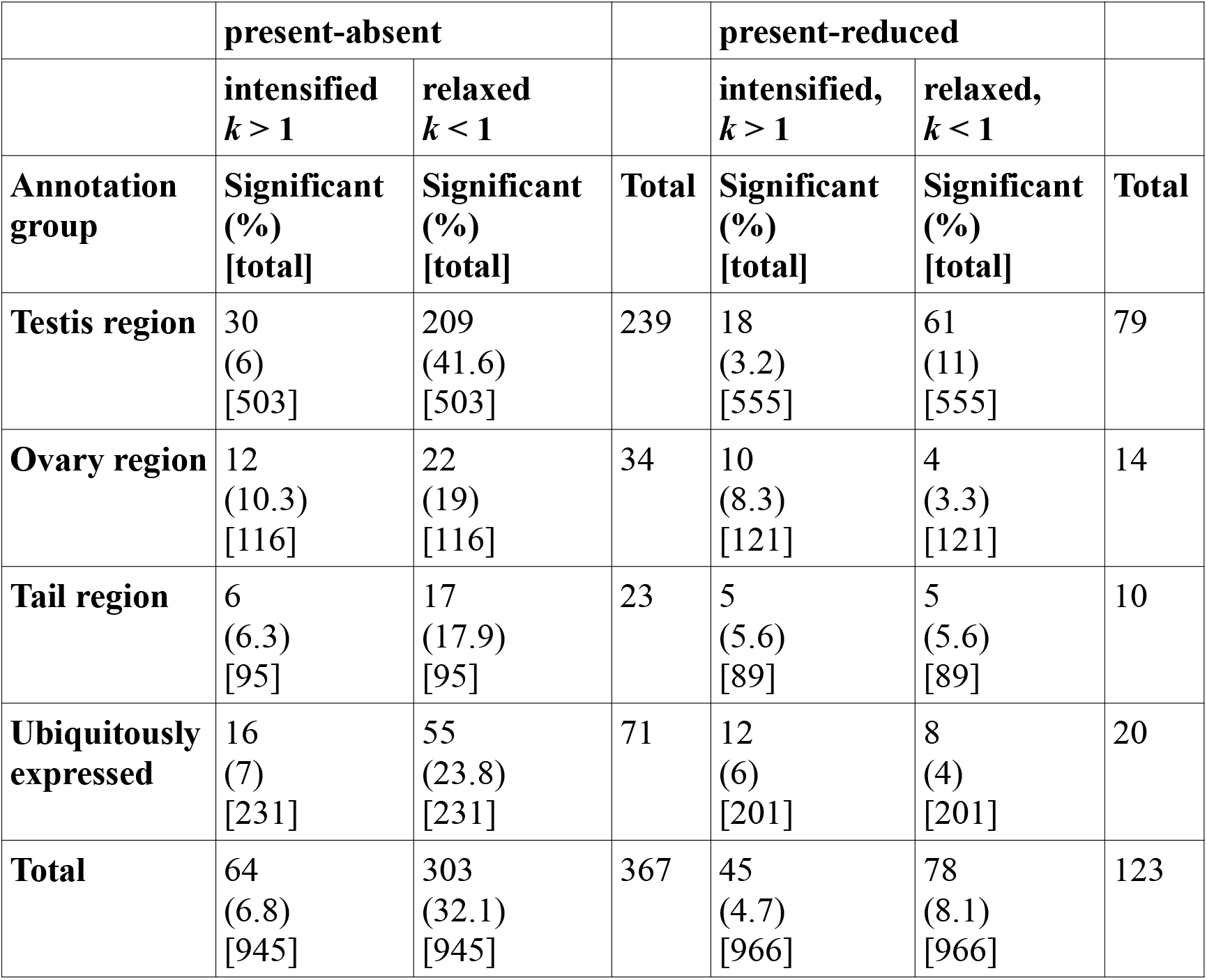
Analyses of molecular sequence evolution using RELAX. The number and percentage of orthogroups (OGs) in each annotation group with a significantly different distribution of ω across sites in test branches (either species with bristles absent or reduced) compared to reference branches (species with bristles present). The percentage of OGs in each annotation group is given in brackets. The total number of OGs tested for each annotation group, from which the expected distribution of counts is computed, is given in square brackets. Results are shown for both contrasts tested: species with bristles present *vs.* species with bristles absent (present-absent), and species with bristles present *vs*. species with reduced bristles (present-reduced). The data are split according to whether the *k* parameter is inferred to be > 1 or < 1.

In the present-reduced contrast, the proportion of OGs with evidence for a different ω distribution among target species was generally lower and showed less variation across annotation groups (table 2). There was a non-significant trend for a deviation from the expected distribution for OGs with *k* > 1 (χ^2^-tests, χ^2^ = 6.7, d.f. = 3, p = 0.08), with an excess of ovary-region (observed, 10; expected, 6) and ubiquitously-expressed (observed, 12; expected, 9) OGs. For OGs with *k* < 1 there was a significant deviation from the expected distribution (χ^2^-tests, χ^2^ = 14.1, d.f. = 3, p < 0.001), with an excess of testis-region OGs (observed, 61; expected, 45), as observed for the present-absent contrast. Moreover, also here there was no significant difference in *k* across annotation groups (Kruskal-Wallis rank-sum tests, *k* > 1, χ^2^ = 1.01, d.f. = 3, p = 0.8; *k* < 1, χ^2^ = 6.1, d.f. = 3, p = 0.11; figure 4B). Figure S11 (with OGs chosen as for figure S10) shows that, just as in figure S9 and S10, in OGs with *k* > 1 most sites were under more intense purifying selection in species with reduced bristles rather than elevated positive selection.

Counter-intuitively, in neither contrast did we find much overlap between OGs that we identified as showing evidence for positive selection in the branch-site analyses and those with a different distribution of ω between test and reference branches in the RELAX analyses (figure 4 and figure S11). Unexpectedly, among the OGs that do overlap, a larger number were identified as being under relaxed rather than intensified selection (present-absent, *k* < 1; N = 19, *k* > 1; N = 9; present-reduced, *k* < 1; N = 7, *k* > 1; N = 5).

## Discussion

We present a comparative study of molecular sequence evolution of reproduction-related and ubiquitously-expressed genes across the genus *Macrostomum,* a group of simultaneously hermaphroditic flatworms that show tremendous diversity in sperm design and other reproductive morphologies (Schärer et al. 2011; Brand et al. *unpublished data*). We characterised the nature of selection operating at reproduction-related and ubiquitously-expressed genes in species with contrasting and convergent sperm morphologies. These sperm morphologies have evolved convergently multiple times and are strongly associated with different mating strategies involving hypodermic insemination or reciprocal copulation that characterise a large part of the sexual selection context of these species (Schärer et al. 2011; Brand et al. *unpublished data*).

Branch models clearly indicate an overall pattern of higher ω values in testis-, ovary-, and tail-region annotated OGs compared to ubiquitously-expressed OGs (figure 2A). This pattern was also largely recovered within lineages with contrasting sperm morphologies, indicating that OGs containing reproduction-related genes show elevated ω; regardless of sperm morphology (figure 2B, figure 3, figure S2). These patterns are consistent with expectations of rapid evolution due to intense selection on reproduction-related genes seen also in other studies (Vacquier and Swanson 2011; Wilburn and Swanson 2016; but see Dapper and Wade 2016, 2020), including a recent small-scale study investigating protein distance and transcript presence/absence in four *Macrostomum* species (Brand et al. 2020). In addition, we found that species lacking sperm bristles, indicative of hypodermic insemination, generally had higher values of ω. Crucially, even the ubiquitously-expressed OGs, with no obvious reproductive function showed elevated w. This pattern (figure 2, figure 3, and figure S2) strongly suggested that general differences in the evolutionary history of species with contrasting sperm morphologies have influenced molecular sequence evolution throughout the genome, and not just at specific subsets of genes. We discuss possible reasons for this observation in more detail below.

In the branch-site models we found relatively little evidence for convergent positive selection at specific amino acid sites in any annotation group. Additionally, although many OGs with evidence for positive selection had sequences from species representing multiple independent losses or reductions of sperm bristles, many only had a single “foreground” lineage driving that result. Such cases do not allow for associating patterns of positive selection with changes in sperm morphology. Evidence of widespread convergent molecular sequence evolution in other systems is mixed. A few examples of convergent amino acid changes in individual genes or groups of genes have been identified in, for example, the Na^+^ channel proteins of electric fishes (Zakon et al. 2006), or certain mitochondrial genes in snakes and agamid lizards (Castoe et al. 2009). However, when investigating genome-wide patterns for convergent evolution to marine environments, Foote et al. (2015) did not find a strong overall signal. Similarly, few of the convergent increases in haemoglobin-oxygen affinity among birds living at high altitudes can be attributed to convergent amino acid changes (Natarjan et al. 2016). Such variation is perhaps expected given that selection on complex traits with multiple genetic underpinnings need not necessarily target variants in the same genes across lineages (Foote et al. 2015; Chikina et al. 2016). Additionally, selection could act on the same genes, but on different sites within those genes, which is a model of molecular sequence evolution not covered by the framework of branch-site models. Consistent with such a model of less strict convergent selection, recent analyses of convergent molecular sequence evolution, using methods that estimate whole-gene evolutionary rates along phylogenetic branches and identifying genes that are strong outliers, found that many genes show a signal of convergent shifts in selective pressure in response to a shift to a marine environment (Chikina et al. 2016), and in parallel adaptation to subterranean lifestyles in mammals (Partha et al. 2019). Understanding the genetic basis for convergent phenotypic evolution and the extent of convergent evolution at the molecular level remains an active field of research (Sackton and Clark 2019).

The finding that ω values in branch tests were higher at reproduction-related genes, even when accounting for the differences between sperm bristle categories, indicated that selection is indeed more intense at these loci. However, the paucity of evidence for convergent selection at specific amino acid sites in the branch-site models suggested that selection is likely acting in different regions of the genes involved. Additionally, some of the alignments shown in figure S8 revealed that uncertain alignments in fragmented regions could potentially confound the signal of positive selection. Alignment uncertainty could reflect biological reality that is not captured by codon models or it could be due to the incomplete nature of the available *de novo* transcriptome assemblies. Genome-guided transcriptome assembly methods could potentially improve transcriptome contiguity and provide for more robust analyses, but currently there is only a single published genome in the genus *Macrostomum* (Wasik et al. 2015; Wudarski et al. 2017).

Because it is not expected that positive selection should be acting across all genes in the genome, the observed genome-wide signal of elevated ω in species with absent bristles spoke to a general feature of the evolutionary history of these species driving patterns of molecular divergence. This raised the possibility that a process such as relaxed selection due to reduced efficiency of purifying selection, rather than positive selection, could be responsible for our observations of genome-wide elevated ω in those species. While relaxed selection could also result from reduced intensity of selection, this would not be expected to affect genome-wide patterns by increasing mean ω throughout, but rather only target specific genes and thus increase the variance in ω. We found that evidence for relaxed selection was common in the present-absent contrast (overall ∼32% of OGs), and that it was more common than intensified purifying or positive selection (overall ∼5% of OGs). Additionally, there was little overlap between OGs identified as being under positive selection in branch-site tests and OGs identified as being under intensified selection in the RELAX analyses (figure 4, figure S11), which could be partly explained by the different assumptions made in PAML and RELAX. For example, branch-site models in PAML do not allow for positive selection in background branches, while the models in RELAX do (Kosakovsky Pond et al. 2011; Wertheim et al. 2015). Furthermore, the assumption, in RELAX, that intensified selection should simultaneously affect sites under positive selection and sites under purifying selection may not be realistic (i.e. the effects may not necessarily be symmetric).

One possible explanation for genome-wide elevated ω in species with absent bristles is a reduced efficiency of natural selection due to smaller effective population size (*N_e_*) in self-fertilising species. A few species of hypodermically inseminating *Macrostomum*, and at least one reciprocally copulating species (namely *M. mirumnovem*), are known to be capable of self-fertilisation (Ramm et al. 2012; Ramm et al. 2015; Giannakara and Ramm 2017; Giannakara and Ramm 2020, Singh et al. 2020). Some hypodermically inseminating species also show no evidence for inbreeding depression, suggesting they could be preferential self-fertilisers (Giannakara and Ramm 2017), while other species seem to show facultative selfing (Ramm et al. 2012; Ramm et al. 2015). Relaxed selection, or more specifically reduced efficiency of selection, due to self-fertilisation is in fact predicted from theoretical models (Glémin 2007; Glémin et al. 2019). Support for this is found in a number of plant systems, but evidence from self-fertilising animals is much rarer (reviewed in Hartfield 2016). Sexual conflict over sex roles could have repeatedly driven the evolution of alternative mating strategies (i.e. hypodermic insemination), and the resulting adaptations (injecting stylets), in turn, may facilitate the evolution of selfing (Ramm et al. 2012) allowing higher rates of selfing that drive down *N_e_*. This increases ω genome-wide *via* the fixation of deleterious mutations as a result of the reduced efficiency of natural selection (Glémin 2007). We find some evidence for this in that a substantial proportion of OGs show evidence of relaxed selection in species with absent bristles.

The above hypothesis makes the further prediction that overall genetic diversity should be lower in hypodermically mating species. To further explore the relationship between a hypodermic mating strategy and *N_e_*, which may contain clues about selfing rates and the efficiency of selection in these species, population genomic data will be necessary. Additionally, more information about the taxonomic distribution of selfing in *Macrostomum* will be invaluable. For example, it is currently unknown whether any of the intermediate species are capable of selfing. Thus, differences in the mating behaviour among species of *Macrostomum* may be shaping genome-wide patterns of molecular sequence evolution, seemingly to a large extent *via* inefficient selection due to lower *N_e_*, likely resulting from higher selfing rates in hypodermically inseminating species. Meanwhile, sexual selection may still be driving the more rapid evolution observed at reproduction-related genes compared to ubiquitously-expressed genes. This combination of selection and drift is an underappreciated point in the genomic consequences of sexual selection. Different evolutionary forces act together to produce the patterns seen in genomic variation. Such combinations of effects make observed patterns difficult to interpret, especially without functional information with which to classify the genes of interest. Functional information for as many genes as possible is invaluable, and models of molecular sequence evolution that explicitly test for the effects of different evolutionary forces have to be deployed in combination to obtain a comprehensive picture.

In this study we re-capitulate the common finding of rapid evolution at reproduction-related genes and show that this is the case also in hermaphroditic animals. This study therefore highlights the importance of considering diverse sexual systems, including hermaphroditism, as much of the theory and empirical work currently remains focused primarily on separate-sexed organisms. Evidence from hermaphroditic organisms, especially animals, is much rarer. In particular, the effects of sex-biased or sex-limited expression of genes on the efficacy of selection, especially with respect to reproduction-related genes (Dapper and Wade 2016; 2020), can likely be avoided, since in simultaneously hermaphroditic animals a gene cannot be strictly sex-limited in expression. Some studies have investigated rates of sequence evolution at reproduction-related genes in self-compatible, hermaphroditic plants where reproduction-related genes are also often found to evolve rapidly, though the mechanism is still debated (Szövényi et al. 2013; Arunkumar et al. 2013; Gossmann et al. 2013; 2016; Harrison et al. 2019; Moyle et al. 2021). Additionally, the opportunity for haploid selection in plants (e.g. in transcriptionally active pollen) further complicates expectations and comparisons to animal systems, while the evidence for haploid selection in animals is limited, given that it is thought that few genes are expressed in the haploid stages of animals (Joseph and Kirkpatrick 2004, but see Alavioon et al. 2017).

The conditions under which genes could experience differences in the efficiency of selection due to the fact that hermaphroditic individuals have dual sex roles remain largely unexplored. For example, it is plausible that genetic variation for sex allocation (i.e. the degree to which a hermaphroditic individual allocates resources to its male or female function) could become linked to sexually antagonistic genetic variation (“intra-individual sexual antagonism”; Abbott 2011), and that this process might precede the evolution of what could be considered proto-sex chromosomes. Such linkage effects might result in genes that are differentially expressed between more “male-biased” *vs.* more “female-biased” individuals, i.e. genotypes that consistently differ in the relative amount of fitness they obtain through one or the other sex function (“effective gender”; *sensu* Lloyd 1980, sometimes also “functional gender”; Charnov 1982; Verdú et al. 2004). In addition, genes involved in sperm competition traits experience less efficient selection, because the genetic variation in these traits that is present within a multiply mated female only represents a fraction of the total genetic variation in the population, especially when female re-mating rates are low (Dapper and Wade 2016; 2020). Intuition would suggest that a reduced efficiency of selection at genes related to sperm competition should be expected also in hermaphrodites. Thus, some caution in interpretation remains warranted when comparing separate-sexed organisms and hermaphrodites, and relaxed selection should at least be considered as a possible explanation for elevated rates of sequence evolution in (some) reproduction-related genes also for simultaneous hermaphrodites. Exploring these ideas will require further development of theory to clarify expectations.

Finally, our results also allow generating candidates for functional investigations on genes that may underlie some of the interesting divergence in morphology observed throughout the genus *Macrostomum*. In the Supplementary Materials (Supplementary Text T3) we highlight a few OGs that we opportunistically selected based on a combination of i) an examination of individual alignments (including alignment quality and the number of independent losses/reductions of sperm bristles represented), ii) evidence of rapid evolution (for example, values of ω above the median for ubiquitously-expressed OGs from branch models), iii) evidence for positive selection (from branch-site models), and/or iv) more detailed expression information, where available. Several of these represent intriguing candidates for the evolution of reproductive traits, including the sperm bristles. Indeed, one already has an established knockdown phenotype, producing aberrant sperm morphology in *M. lignano* (Grudniewska et al. 2018). Another candidate is known to be expressed in the developing stylet (Weber et al. 2018), which is another aspect of the extreme diversity in reproductive morphology in this genus. Thus, our findings, combined with an examination of the alignments, allow the generation of promising individual candidate genes for further investigation. Some of these genes might be good candidates to explore in the context of differences among species in sperm morphology and other sperm traits, in the stylet morphology, in the seminal fluid, and in other reproductive functions. Combined with more detailed information about gene expression patterns, these candidates can provide great insights and guide the exploration of gene-phenotype associations across the genus.

It is important to note, that our results are mostly based on *de novo* transcriptome assemblies from, typically wild-caught, single individuals. In addition to errors of orthology inference due to ancient or recent whole-genome duplication events, which are known from several species of *Macrostomum* (Zadesenets et al. 2016; 2017a,b; 2020), the orthology detection approach we use here has several heuristic decision steps that may introduce biases. Specifically, assemblies generated from outbred individuals may increase the redundancy of the dataset as a result of isoform and allelic variation. In the face of lineage specific duplications, the approach we used picks the sequence with the most aligned columns (Yang and Smith 2014). Nevertheless, tree-based orthology detection approaches are best placed to at least disambiguate orthology relationships in the face of older duplication events (Yang and Smith 2014). Additionally, we note that reproduction-related genes are here defined on the basis of gene expression patterns in only a single species, namely *M. lignano* (Arbore et al. 2015; Brand et al. 2020). It therefore remains possible that gene expression patterns change and that genes either gain or lose reproduction-related functions as species diverge. Further gene expression studies from other species, which are currently under way, would greatly increase the reliability with which we can classify genes. Finally, strong sexual selection on reproduction-related genes should also leave a population genetic signature detectable in molecular diversity data from within-species datasets (Dapper and Wade 2020). Such datasets do not yet exist for *Macrostomum*, but would clearly be valuable additions. Despite the above caveats, our data and findings represent a significant addition to the understanding of molecular evolution at reproduction related genes and generate further hypotheses about the consequences of sexual selection in the genus *Macrostomum*.

## Conclusions

We have conducted a comparative study of molecular sequence evolution in a genus of simultaneously hermaphroditic flatworms, which, to our knowledge, is the first robust study of molecular sequence evolution at reproduction-related genes in hermaphroditic animals. This study highlights the importance of studies that contrast lineages with different mating strategies and associated morphologies and behaviours, which presumably also differ in the sexual selection context they experience. Importantly, the extent of taxon sampling allows the comparison of multiple independent changes in sperm morphology (and the associated mating strategies), a feature largely lacking from previous studies. Furthermore, we expand the scope of work investigating the effects of sexual selection on genomic patterns of variation to include hermaphroditic animals. We show that changes in sperm morphology are associated with elevated ω values throughout the genome, which suggests a general feature of the evolutionary history of species lacking bristles driving patterns of sequence divergence. A possible explanation lies in the mating strategy associated with the bristle-less sperm morphology, namely hypodermic insemination. Given what we have outlined above, we speculate that hypodermically inseminating species may engage in higher rates of self-fertilisation, which should drive down effective population sizes and increase the effect of genetic drift throughout the genome. On top of these genome-wide patterns we also find higher ω values at reproduction-related genes consistent with a strong effect of sexual selection acting on such genes.

## Materials and Methods

### Candidate reproduction-related genes

Previous work on the model species *M. lignano* has characterised gene expression in different body regions of adult worms using a “positional RNA-Seq” approach (Arbore et al. 2015). Briefly, many individual worms were either cut i) once just anterior to the testes, ii) between testes and ovaries, iii) just posterior to the ovaries, or iv) not at all, yielding four types of RNA samples that represented pools of progressively larger fragments of the worm. To make use of more recent and improved genome and transcriptome assemblies (Wudarski et al. 2017; Grudniewska et al. 2018), Brand et al. (2020) repeated the analyses of that RNA-Seq data using transcriptome assembly Mlig_RNA_3_7_DV1.v3 (Grudniewska et al. 2018). By comparing gene expression of genes between neighbouring worm fragments, transcripts were identified with specific expression in the testis-, ovary-, or tail-region of the worm, while other transcripts could be identified as being likely ubiquitously-expressed across the regions (figure S12). Additionally, some transcripts were annotated as being “low-expressed” (i.e. with <50 reads mapping from the uncut worm sample) for downstream filtering (see Additional file 14 in Brand et al. 2020).

### Ortholog inference and codon-alignment preparation

We made use of recently generated whole transcriptome data for 97 species of *Macrostomum* and three outgroups (i.e. three undescribed species of the sister genus *Psammomacrostomum*; Janssen et al. 2015), including clusters of homologous sequences from across all species (for details see Brand et al. *in press*). Briefly, homolog groups for all identified protein-coding genes in the assembled transcriptomes were obtained by first conducting an all-vs-all BLASTp search and then clustering these results using OrthoFinder (v. 1.1.10; Emms and Kelly 2015), keeping only homolog groups containing at least 10 species. These homolog groups were aligned with MAFFT (v. 7.3; Katoh and Standley 2013), the alignments were trimmed with ZORRO (v. 1.0; Wu et al. 2012), and finally gene trees were built with IQ-TREE (v. 1.5.6; Nguyen et al. 2015) using the best fitting model of molecular sequence evolution for each alignment (Brand et al. *in press*).

For our study, we identified those homolog groups, and their associated gene trees, which contained a *M. lignano* gene, either from one of the above-mentioned reproduction-related tissue regions or from a randomly chosen set of 1,000 ubiquitously-expressed genes, and annotated them according to the inferred function of the *M. lignano* gene (yielding four functional annotation groups). We then pruned gene trees in three steps according to the procedure outlined by Yang and Smith (2014). First, we pruned tips from the tree if they were longer than an absolute length of 2 substitutions per site or if the length of a tip was > 10x the length of its sister tip. Next, if a clade was composed of multiple sequences from just a single species, we retained only the sequence with the most aligned columns. Finally, we split trees into subtrees at long internal branches, and only retained the resulting subtrees if the number of species in these trees was ≥ 5. Finally, we inferred ortholog groups (OGs) from the resulting subtrees using both the “rooted ingroups” (RT) and “maximum inclusion” (MI) algorithms (Yang and Smith 2014). We used the union of OGs inferred by both of these methods in downstream analyses, always keeping the OG produced by RT where there was overlap, because they were typically larger.

We then extracted the amino acid and nucleotide sequences corresponding to the coding region of these OGs from each species’ transcriptome assembly, aligned them using MAFFT, and converted the amino acid alignments to codon alignments with pal2nal (v. 14; Suyama et al. 2006). We subsequently trimmed these alignments by first computing column-wise alignment scores for the amino acid alignments with ZORRO and then removing the codon columns from the codon alignments that corresponded to low-scoring (score ≤ 3) amino acid columns. If trimmed, gap-less sequences were shorter than 50% of the alignment length, we removed these sequences from the alignment. Finally, we checked alignments for internal stop codons and removed alignments < 80 nucleotides long. Note that these last alignment trimming steps in some cases led to the removal of the original *M. lignano* sequence from the alignment. Where this occurred (N = 19, 1.7%), we nevertheless retained these OGs, preserving the annotation from *M. lignano*, for further analysis. We used a recently produced maximum likelihood species tree based on 385 genes with an occupancy rate of >80% across the genus *Macrostomum*, representing 94,625 aligned amino acids (and referred to as “H-IQ-TREE” in Brand et al. *in press*; figure 1C). Since our alignments contained different sets and numbers of species, we pruned the species tree to contain only the species actually present in each alignment. Only alignments with at least four species were retained.

The pipeline outlined above involves many steps, each of which has the potential to discard genes from the analysis. To evaluate the success of recovering OGs from across the species, we quantified the proportion of OGs remaining that contained an annotated gene from *M. lignano* after each step (figure S1). We found that there is a substantial loss during orthology identification regardless of the functional annotation of the *M. lignano* gene (table S3; figure S1). However, the trend suggests that recovery is somewhat lower (13.6-16.4%) for reproduction-related genes than for ubiquitously expressed genes (25.7%). These results suggest that ortholog identification, in general, has a low recovery rate, at least on the phylogenetic scales investigated here, but also that recovery is worse for groups of reproduction-related genes that are expected to be evolving more rapidly (see also Brand et al. 2020). For all analyses below, the final trimmed alignments used are made available in a supplementary archive (Supplementary Archive 1, available from: https://doi.org/10.5281/zenodo.4972188).

### Branch models

Brand et al. (*unpublished data*) collected morphological data from species across the genus *Macrostomum* (presented in Tables S2 and S3 of that study). In order to test whether species with different sperm morphologies show evidence for different patterns of molecular sequence evolution we first classified species into three categories of sperm morphology according to Brand et al. (*unpublished data*; figure 1B), namely species with bristles present, reduced, or absent. We then identified orthologs that showed variation in *dN/dS* ratios across these three bristle states by fitting a series of codon substitution branch models using the codeml software in the PAML package (v.4.9; Yang et al. 2000; Yang 2007). As outlined in the Introduction, *dN/dS* (ω) is the ratio of non-synonymous (*dN*) to synonymous (*dS*) substitution rates. This value allows inference of selection on a protein-coding gene, with ω = 1 implying that the gene is evolving neutrally, while ω > 1 or ω < 1 are generally considered to imply positive or purifying selection, respectively (but see also below). First, we fit a model that assumes a single ω across the entire tree for each OG alignment. Second, we fit a model where terminal branches leading to species with reduced bristles and species with absent bristles were labelled separately, effectively assuming a different ω for each of the three bristle states (note that the alternative model does not require that all branches have different estimates for ω). Both models have the parameters “fix_omega = 0”, “omega = 0.001, “fix_kappa = 0”, “kappa = 2”, differing only in the input tree with either a single branch label (null model) or three branch labels (alternative model). These models are then compared using a likelihood-ratio test (LRT) with 2 degrees of freedom for each model comparison. FDR was controlled at a rate of 0.05 by converting p-values from the LRT to q-values (Storey and Tibshirani 2003). Note that, by definition, q-values can never be larger than the estimated π_0_ parameter, and therefore with a low estimate for π_0_, q-values can be smaller than the corresponding p-value. We then compare ω estimates from the model with a single ω between annotation groups using a Kruskal-Wallis rank-sum test, followed by a Dwass-Steele-Critchlow-Fligner all-pairs post-hoc test. We also tested whether the proportion of OGs for which the alternative model was a better fit, was larger than expected in some annotation groups using χ^2^-tests. The expected proportions were based on the distribution of all OGs across annotation groups.

Finally, we evaluated the mean level of codon usage bias for species without bristles using the method of Karlin et al. (2001), always setting species with bristles in each alignment as the reference set. This approach determines whether there are codon-usage biases among species with absent bristles compared to species with bristles present. Such a systematic difference could potentially be a confounding variable in a comparison of molecular sequence evolution between two groups.

### Branch-site models

The branch models we described above fit average estimates of ω across the entire alignment, but the entire coding sequence of a gene is often not expected to be under the same kind of selection. Rather, in many cases only a few amino acid sites are likely the targets of intense selection (Yang and Nielsen 2002; Zhang et al. 2005). Therefore, we also fit branch-site models, as implemented in PAML. Since branch-site models only allow for two branch classes (“foreground” and “background”), we performed two contrasts. We set species with absent bristles or species with reduced bristles as the foreground branch in the present-absent and the present-reduced contrasts, respectively, and species with bristles present were always set as the background branches. We chose these contrasts because reciprocal copulation is thought to be the ancestral state for a large part of the *Macrostomum* phylogeny (Brand et al. *unpublished data*).

We fit the branch-site model A (Zhang et al. 2005), which has four site classes (0, 1, 2a, and 2b). This model allows two of these classes to have ω > 1 among branches labelled as foreground (namely 2a and 2b, with parameter ω_2_). Model A was specified with the following PAML parameters “model = 2”, “NSsites = 2”, “fix_omega = 0”, “omega = 0.001”. The other site classes are constrained to 0 < ω < 1 (site class 0, with parameter ω_0_), or ω = 1 (site class 1, with parameter ω_1_) in both background and foreground branches. We also fit the corresponding null model (A_null_, specified with PAML parameters “model = 2”, “NSsites = 2”, “fix_omega = 1”, “omega = 1”). A_null_ is otherwise identical, but the parameter ω_2_ is fixed to 1 (Zhang et al. 2005). Thus, the null model allows for sites that were under purifying selection in the background branches, but are now evolving neutrally in the foreground branches (site class 2a) as well as sites that are evolving neutrally in both the background and foreground branches (site class 2b; see Zhang et al. 2005 for full details). The null and alternative models were then compared using an LRT with 2 degrees of freedom (as above). Branch-site tests, as implemented in PAML, also allow the assignment to each site in the alignment of a posterior probability that it belongs to a particular site class (Yang et al. 2005). The probability that a site belongs to a site class under positive selection in the foreground branch is of particular interest. Thus, for those OGs with evidence of positive selection we identified sites with a high posterior probability (>0.95) of being under positive selection.

These branch-site tests amount to a stringent test for a pattern of positive selection, in common to the lineages either lacking or having reduced sperm bristles, on a few sites of a protein-coding gene. Where alignments include multiple independent origins (see also figure S5), this can be taken as a test of convergent molecular sequence evolution associated with shifts in sperm morphology. Though many OGs contain multiple species with absent bristles, these may often only represent one or two phylogenetically independent losses (e.g. multiple representatives of the morphologically uniform hypodermically mating clade, top of figure 1C). OGs with more independent changes are arguably more informative concerning convergent molecular sequence evolution, and therefore more informative about patterns of molecular sequence evolution associated with differences or changes in bristle morphology. Simply correcting for multiple testing would not account for the variation across OGs in the number of phylogenetically independent losses or reductions of bristles that are represented within each OG. Therefore, in order to keep as much information from independent changes as possible, we chose to retain all OGs with an uncorrected LRT p-value < 0.05. This permitted a more nuanced exploration of results stratified by the number of independent losses within OGs. However, we nevertheless also provide overall q-values (as for the branch tests above). Among genes inferred to show evidence for positive selection (LRT p-values < 0.05) we tested whether there were differences in the ω_2_ parameter estimates from foreground branches across annotation groups using Kruskal-Wallis rank-sum tests. We further tested whether some annotation groups had a larger proportion of OGs for which the alternative model was a better fit, than expected based on the distribution of OGs tested as a proportion of all OGs using χ^2^-tests.

### RELAX models

Finally, we also performed analyses with RELAX (Wertheim et al. 2015) in the HyPhy suite of analysis tools (v. 2.5.29; Kosakovsky Pond et al. 2019). In the hypothesis-testing framework we use here, RELAX compares a set of test branches to a set of reference branches in a phylogeny, in order to test if the distribution of ω values across sites is different between them. Specifically, it fits a discrete distribution of ω values with *n =* 3 site categories to model the distribution of ω (such that the three categories have ω_0_ ≤ ω_1_ ≤ 1 ≤ ω_2_), and the proportion of sites in each category is also estimated from the data. It further assumes that the ω values of the distribution for the test branches is related to the distribution in reference branches by a value, *k,* that describes either relaxation or intensification of selection. It does so by comparing two models by an LRT, namely a null model where *k* = 1 and an alternative model where *k* can take any value > 0. If the LRT rejects the null model it indicates that selection is either intensified (when *k* > 1) or that selection is relaxed (when *k* < 1). Note that in the hypothesis-testing framework, the proportion of sites in each category is constrained to be the same across test and reference branches. RELAX can also fit a less constrained Partitioned Descriptive Model (PDM) where the ω values as well as the proportion of sites in each category may vary between test and reference branches. The PDM estimates can then be used to make more refined interpretations of differences between test and reference branches, while hypothesis testing is performed with the null and alternative models described above (Wertheim et al. 2015).

In the RELAX framework, relaxation of selection should result in the distribution of ω values becoming more narrowly distributed around a value of ω = 1 (Wertheim et al. 2015), since *dN* and *dS* become more similar. In contrast, intensified selection results in the distribution of ω becoming wider. In other words, RELAX assumes that intensified selection simultaneously intensifies both positive and purifying selection throughout the gene. As above, we ran RELAX for data sets that only include two sperm bristle categories (present-absent and present-reduced), and we adjusted LRT p-values with the q-value method to control the FDR. We then split those data where the null model was rejected into those where *k* < 1 and *k* > 1, respectively. Finally, we compared the distribution of *k* values across the reproduction-related annotation groups within these two groups using Kruskal-Wallis rank-sum tests.

### Data analysis and processing

Unless otherwise stated, all statistical tests have been performed in R (v. 3.6.3; R Development Core Team 2020) and figures are plotted with the “ggplot2” package (v. 3.3.1; Wickham 2016) with the aid of various helper packages: “reshape” (v. 0.8.8; Wickham 2007), “plyr” (v. 1.8.6; Wickham 2011), and “gridExtra” (v. 2.3; Auguie 2017). The Dwass-Steele-Critchlow-Fligner post-hoc all-pairs tests were performed with tools from the “PCMRplus” package (v. 1.4.4; Pohlert 2020), MANOVA and mixed model results given in the Supplementary Materials were obtained with the “MANOVA.RM” (v. 0.4.3; Friedrich et al. 2021) and “nlme” (v. 3.1-149; Pinheiro et al. 2020) R packages respectively. Q-values were computed using the “qvalue” package (v. 2.16; Storey et al. 2019). The phylogenetic tree was processed and plotted with tools from the “ape” (v. 5.4; Paradis and Schliep 2019), “phytools” (v. 0.7-47; Revell 2012), and “ggtree” (v. 1.17.4; Yu et al. 2017) packages. Codon usage bias analyses were performed with the “coRdon” package (v. 1.2.0; Elek et al. 2019). See the R scripts in the supplementary materials for full details.

## Acknowledgments

This work was supported by grants from the Swiss National Science Foundation (grant numbers 31003A_162543 and 310030_184916 to LS). We also thank Lukas Zimmermann for IT support. We would also like to thank Darren Parker, Mike Wade, and Lynda Delph for valuable discussion. Computations were performed, in part, at sciCORE (http://scicore.unibas.ch/) scientific computing center at the University of Basel.

## Data availability

Transcriptome assemblies forming the basis of these analyses are available from https://doi.org/10.5281/zenodo.4543289. These transcriptome assemblies are themselves based on raw sequencing data which has been deposited on NCBI: [BioProject accession: PRJNA635941]. The final codon alignments, in addition to analysis scripts and datasets used in the above analyses are provided as supplementary files to this manuscript and are available from: https://doi.org/10.5281/zenodo.4972188.

