## Supplementary material for "Faster rates of molecular sequence evolution in reproduction-related genes and in species with hypodermic sperm morphologies": Figure S8

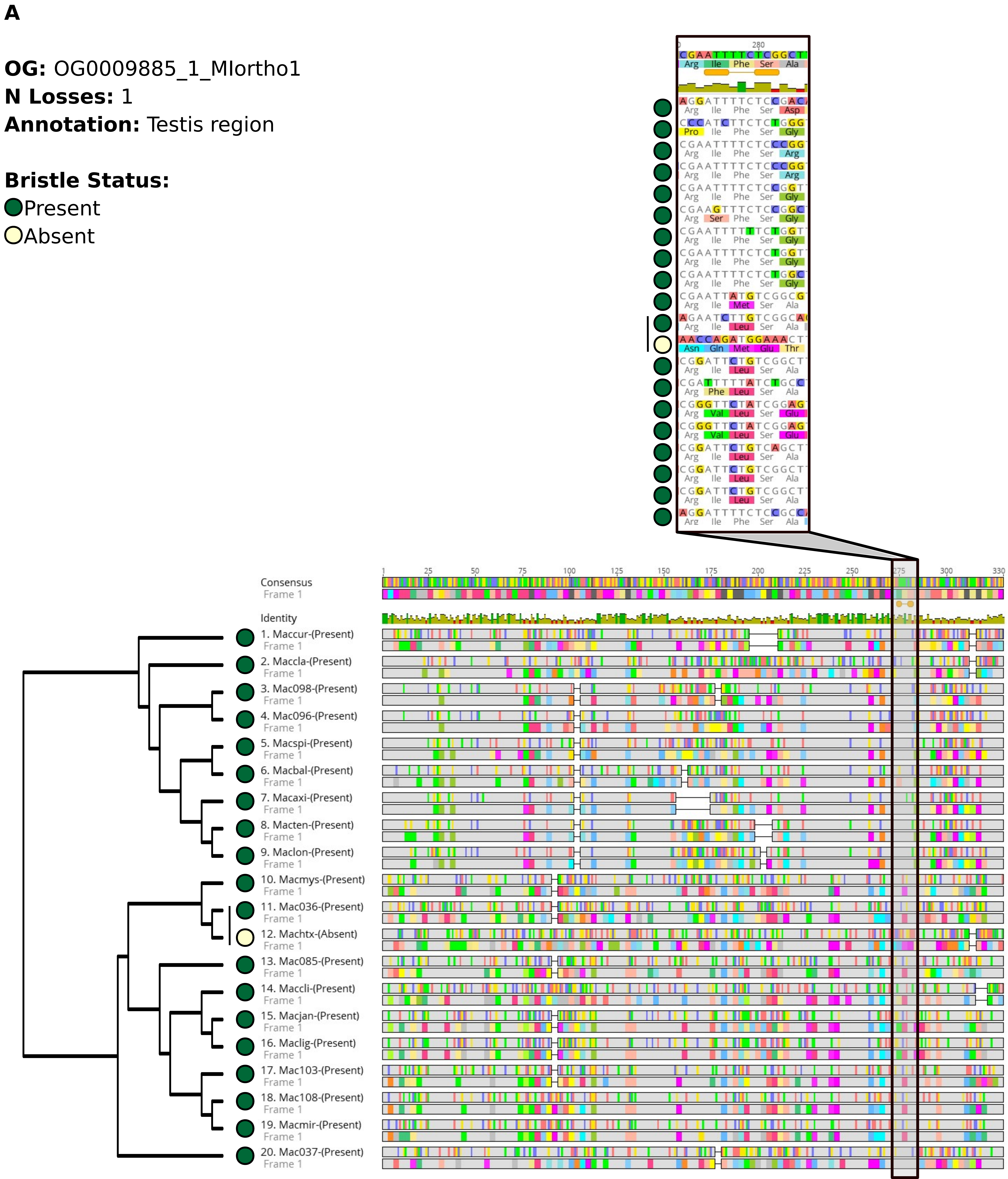
A
OG: OG0009885_1_MIortho1
N Losses: 1
Annotation: Testis region
Bristle Status:
Present
Absent
Consensus
Frame 1
Identity
1. Maccur-(Present)
2. Maccla-(Present)
3. Mac098-(Present)
4. Mac096-(Present)
5. Macspi-(Present)
6. Macbal-(Present)
7. Macaxi-(Present)
8. Macten-(Present)
9. Maclon-(Present)
10. Macmys-(Present)
11. Mac036-(Present)
12. Machtx-(Absent)
13. Mac085-(Present)
14. Maccli-(Present)
15. Macjan-(Present)
16. Maclig-(Present)
17. Mac103-(Present)
18. Mac108-(Present)
19. Macmir-(Present)
20. Mac037-(Present)

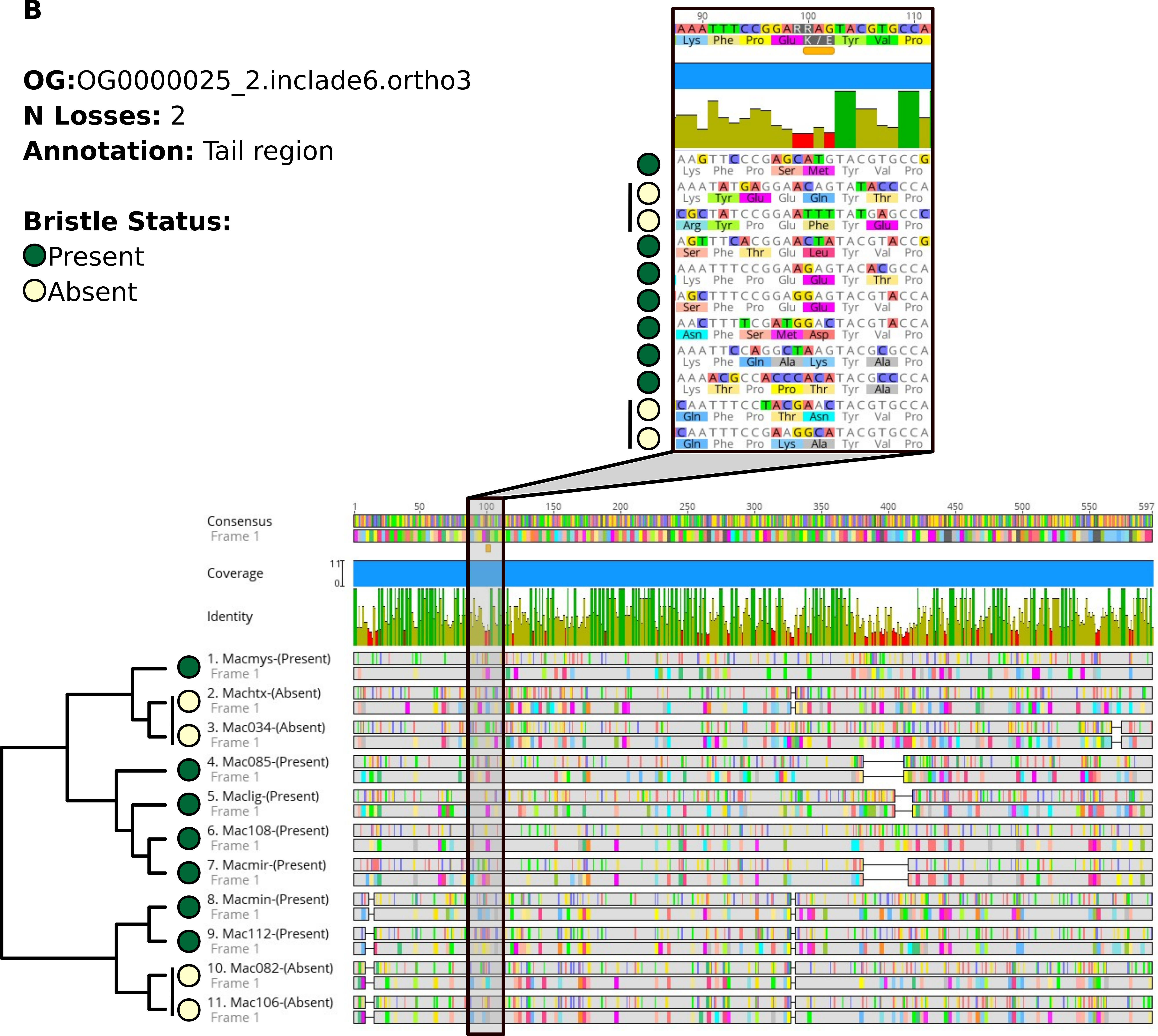
B
OG:OG0000025_2.inclade6.ortho3
N Losses: 2
Annotation: Tail region
Bristle Status:
Present
Absent
Consensus
Frame 1
Coverage
Identity
1. Macmys-(Present)
2. Machtx-(Absent)
3. Mac034-(Absent)
4. Mac085-(Present)
5. Maclig-(Present)
6. Mac108-(Present)
7. Macmir-(Present)
8. Macmin-(Present)
9. Mac112-(Present)
10. Mac082-(Absent)
11. Mac106-(Absent)

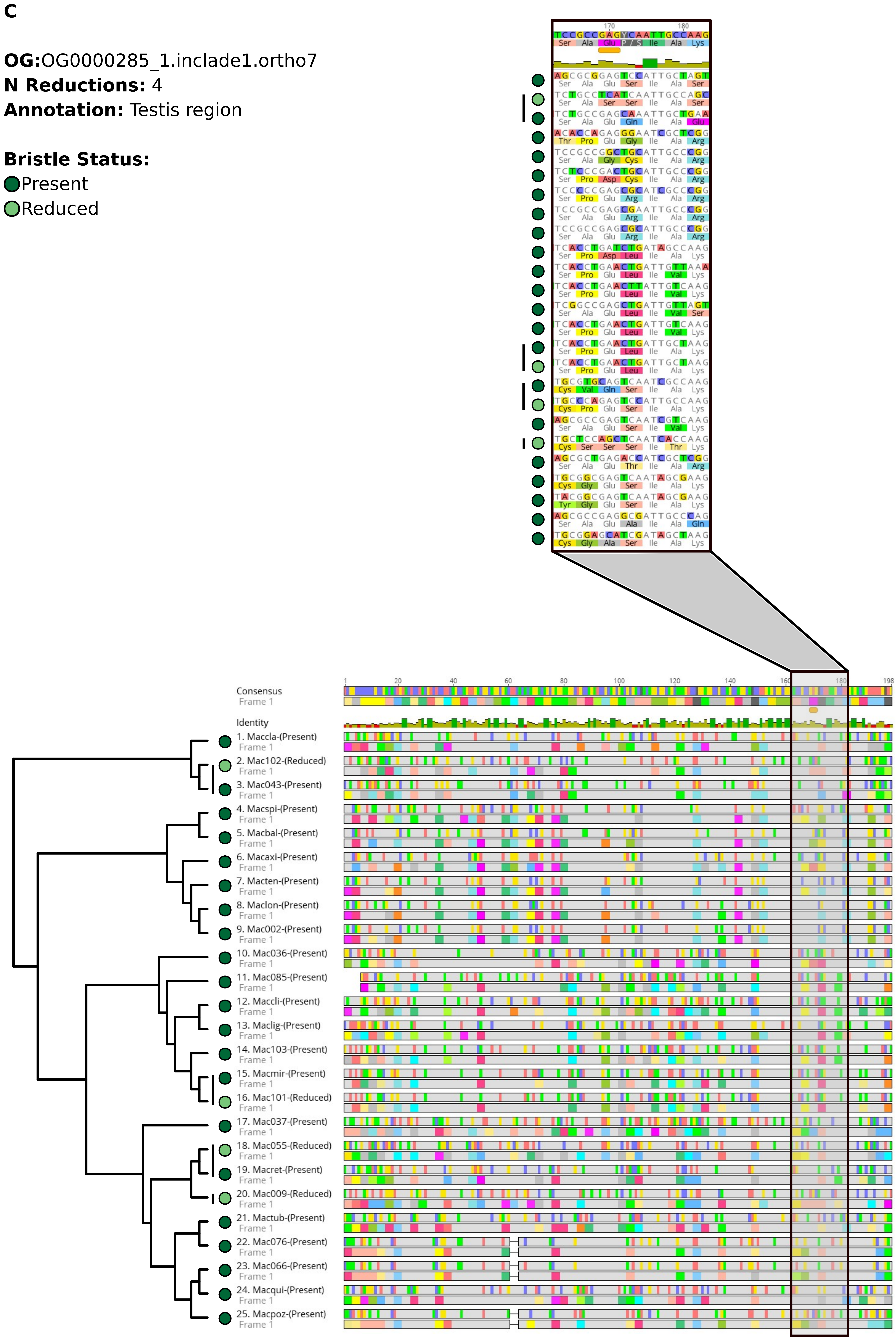
C
OG:OG0000285_1.inclade1.ortho7
N Reductions: 4
Annotation: Testis region
Bristle Status:
Present
Reduced
Consensus
Frame 1
Identity
1. Maccla-(Present)
2. Mac102-(Reduced)
3. Mac043-(Present)
4. Macspi-(Present)
5. Macbal-(Present)
6. Macaxi-(Present)
7. Macten-(Present)
8. Maclon-(Present)
9. Mac002-(Present)
10. Mac036-(Present)
11. Mac085-(Present)
12. Maccli-(Present)
13. Maclig-(Present)
14. Mac103-(Present)
15. Macmir-(Present)
16. Mac101-(Reduced)
17. Mac037-(Present)
18. Mac055-(Reduced)
19. Macret-(Present)
20. Mac009-(Reduced)
21. Mactub-(Present)
22. Mac076-(Present)
23. Mac066-(Present)
24. Macqui-(Present)
25. Macpoz-(Present)

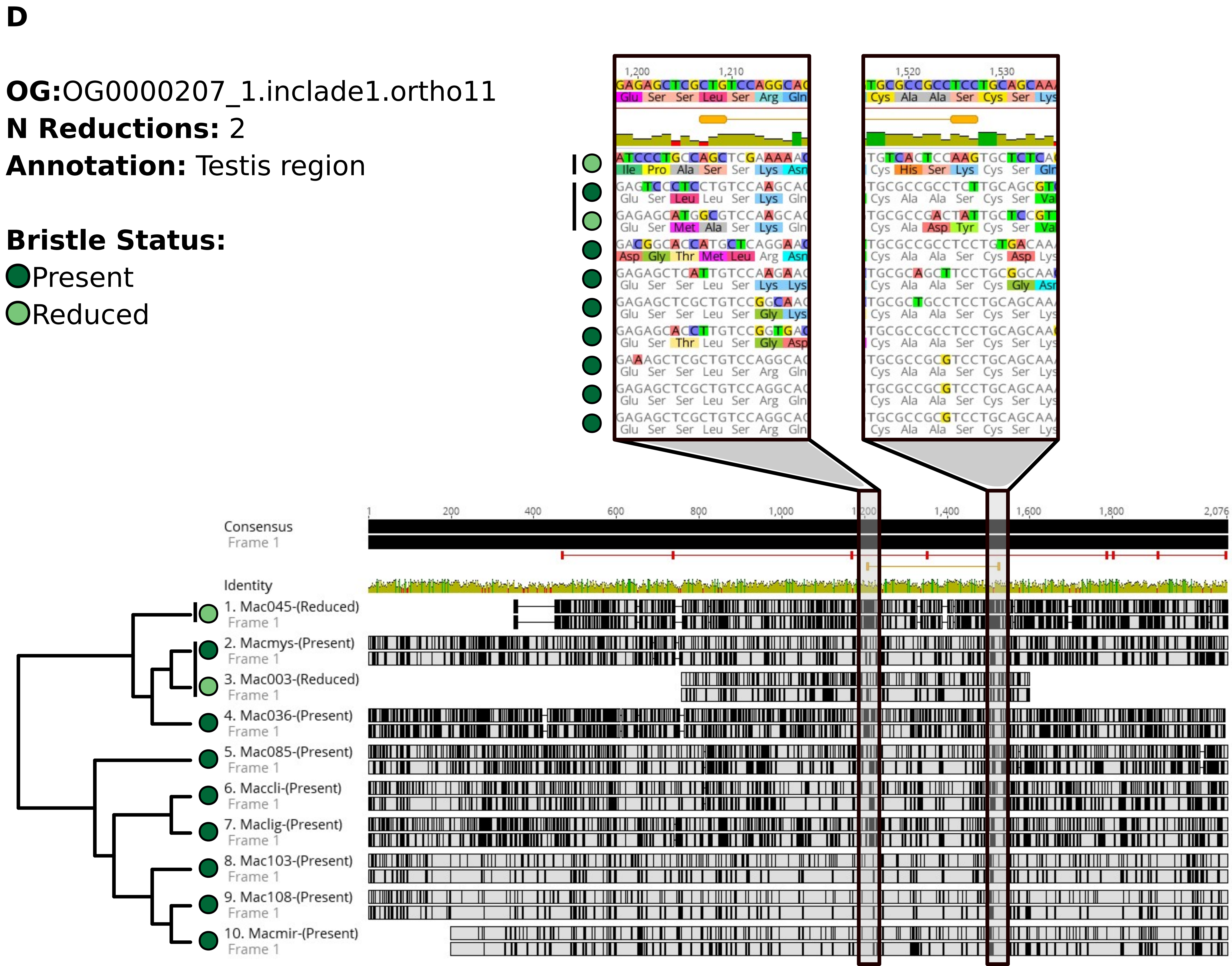
D
OG:OG0000207_1.inclade1.ortho11
N Reductions: 2
Annotation: Testis region
Bristle Status:
Present
Reduced
Consensus
Frame 1
Identity
1. Mac045-(Reduced)
2. Macmys-(Present)
3. Mac003-(Reduced)
4. Mac036-(Present)
5. Mac085-(Present)
6. Maccli-(Present)
7. Maclig-(Present)
8. Mac103-(Present)
9. Mac108-(Present)
10. Macmir-(Present)

**E**

**OG:**OG0000247_1.inclade3.ortho5

**N Losses:** 3

**Annotation:** Testis region

**Bristle Status:**

● Present

○ Absent

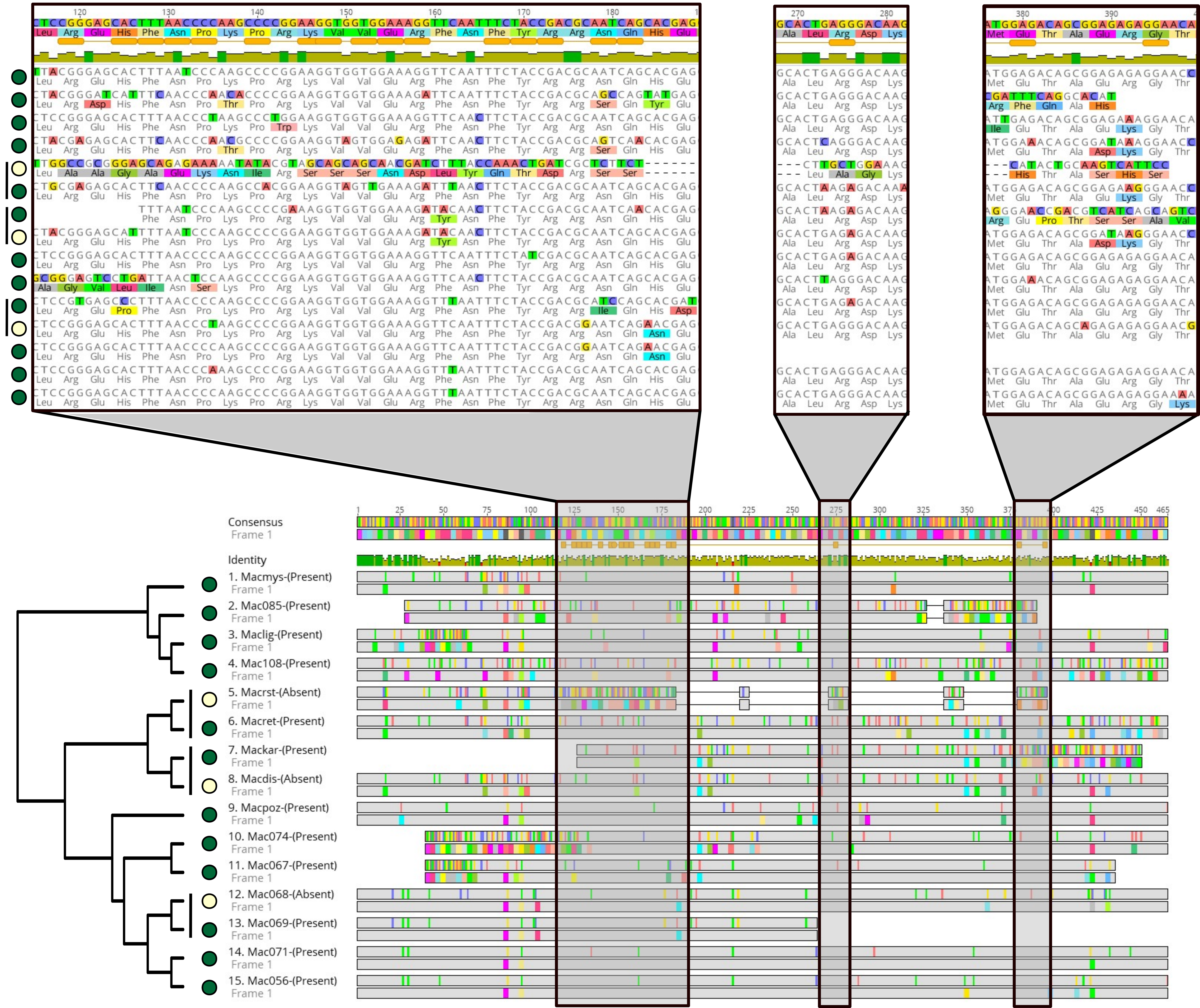

**F**

**OG:**OG0001675_2_MIortho3

**N Losses:** 1

**Annotation:** Ovary region

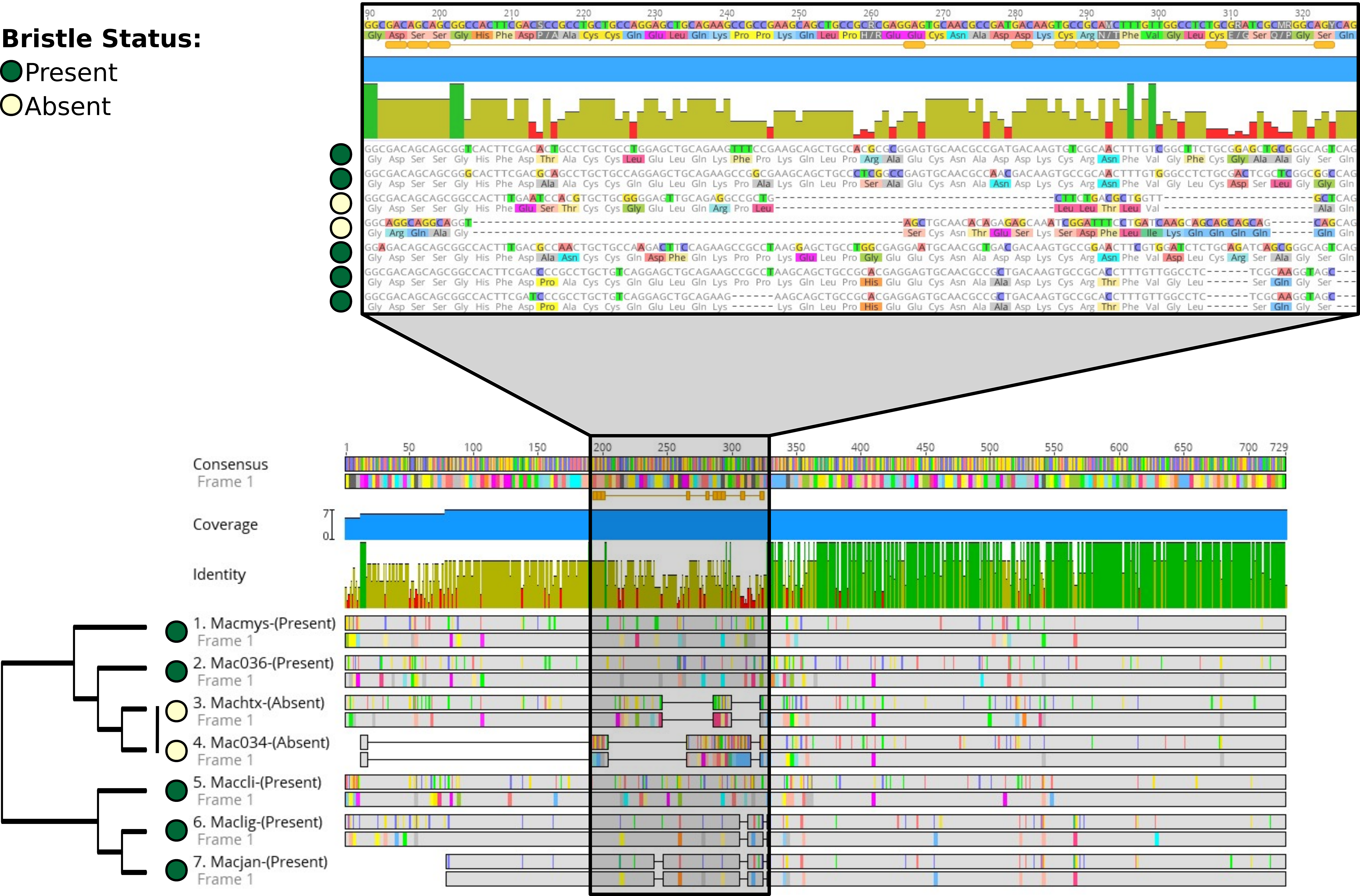

**G**

**OG:**OG0000113_4.inclade1.ortho5

**N Reductions:** 1

**Annotation:** Ubiquitously expre

**Bristle Status:**

Present

Reduced

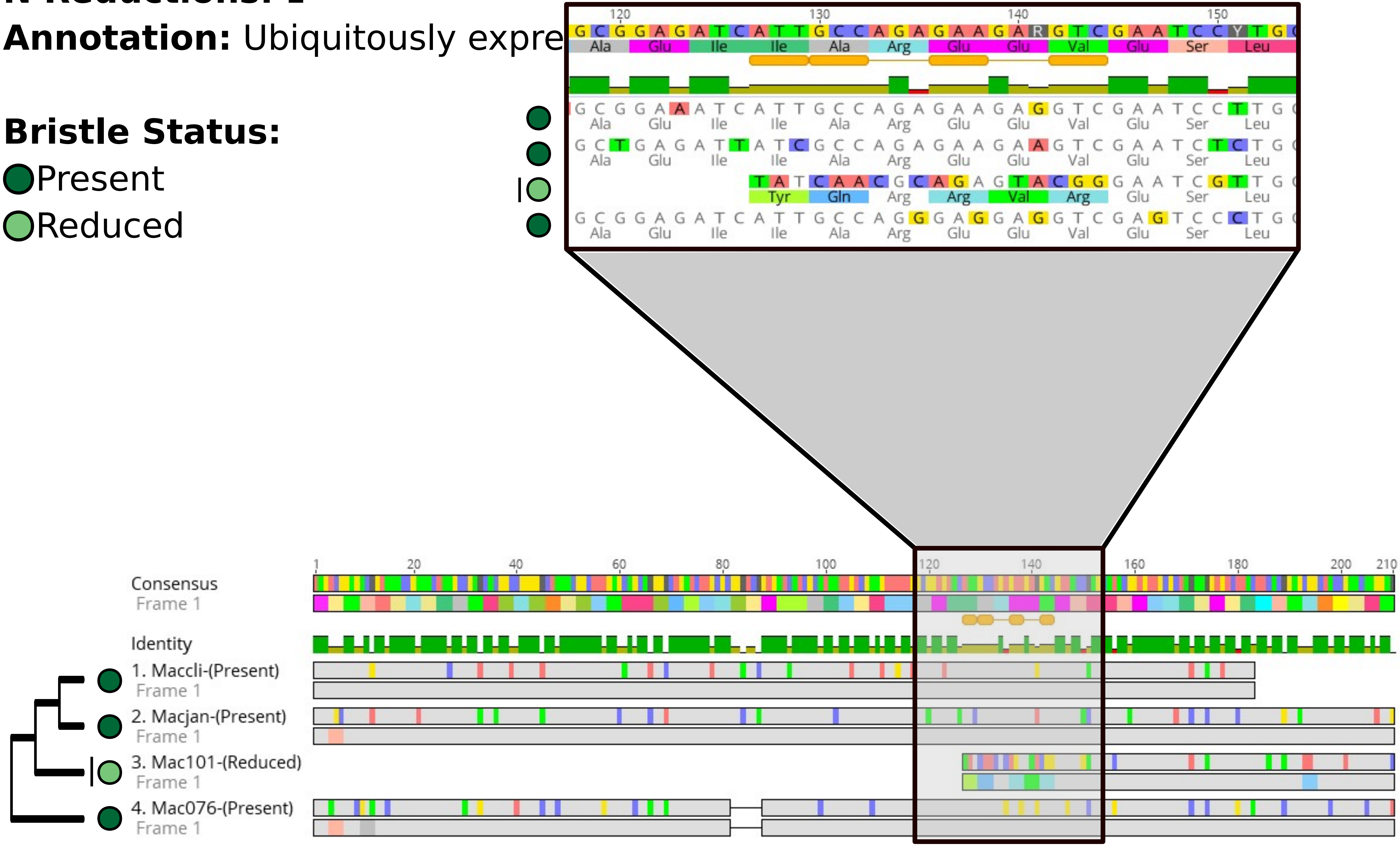

**H**

**OG:**OG0011654_1_MIortho1

**N Losses:** 5

**Annotation:** Ovary region

**Bristle Status:**

Present

Absent

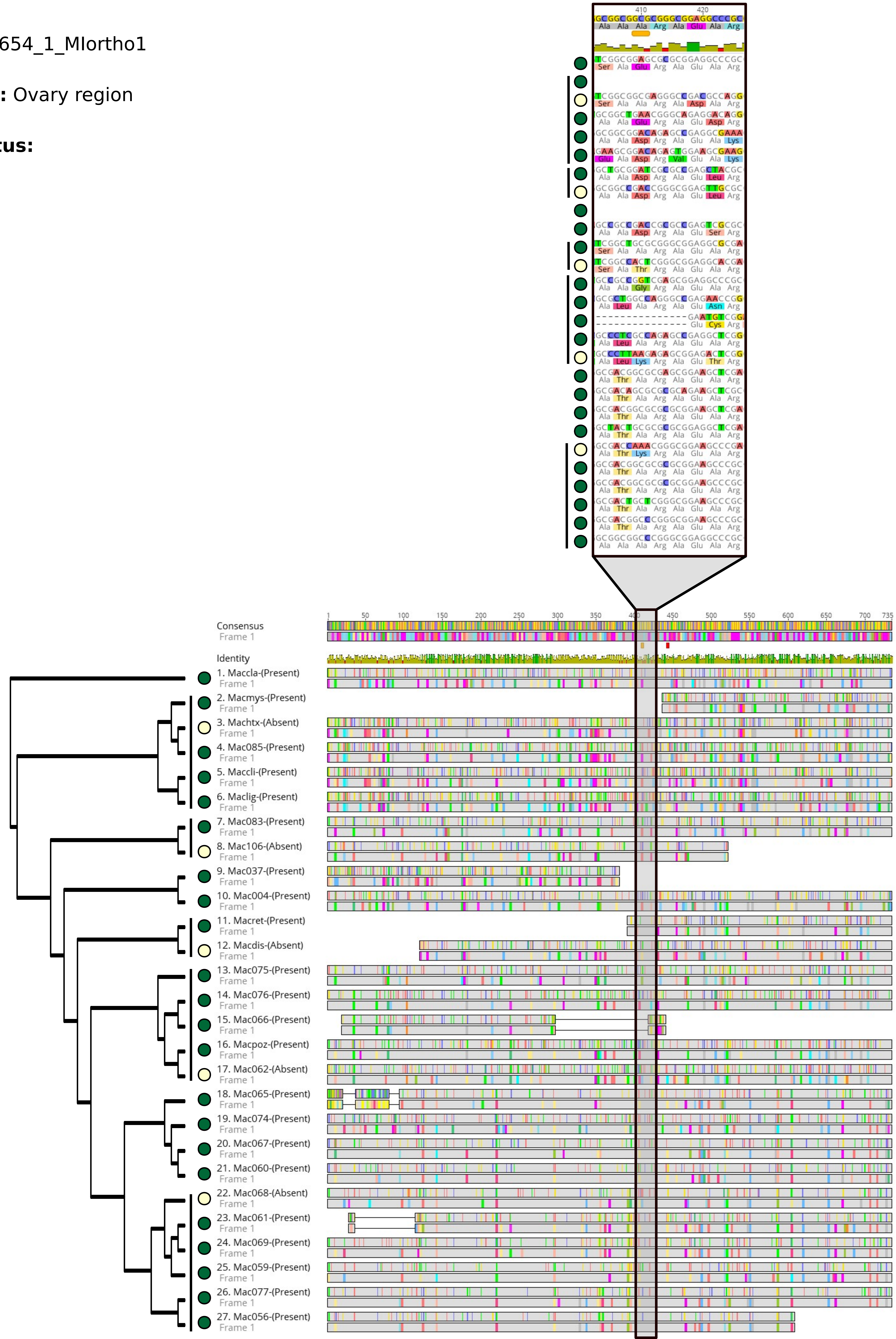

**I**

**OG:**OG0000222_1.inclade1.ortho11

**N Losses:** 2

**Annotation:** Ovary region

**Bristle Status:**

Present

Absent

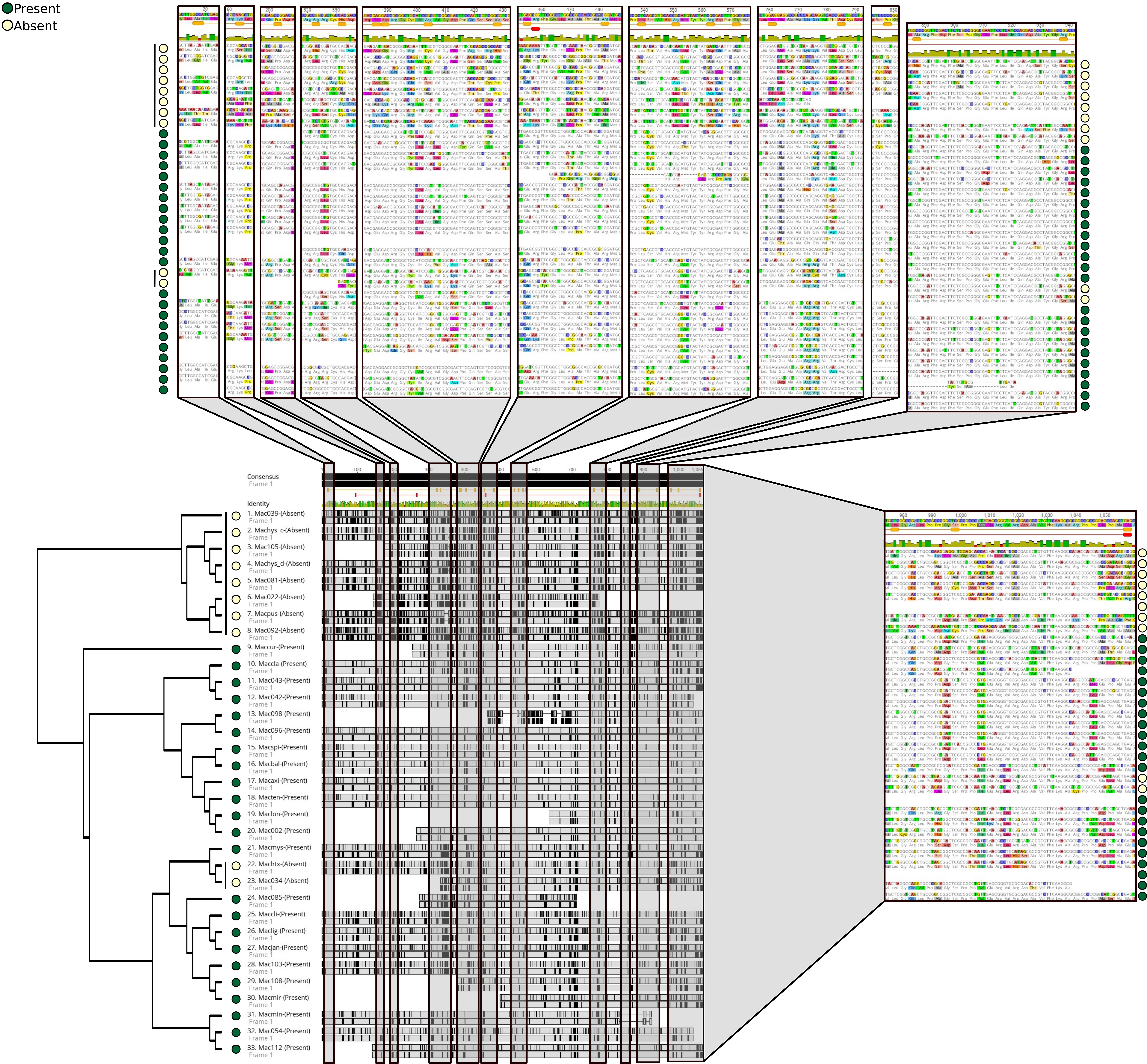

**J**

**OG:**OG0003507_1.inclade1.ortho3

**N Reductions:** 2

**Annotation:** Tail region

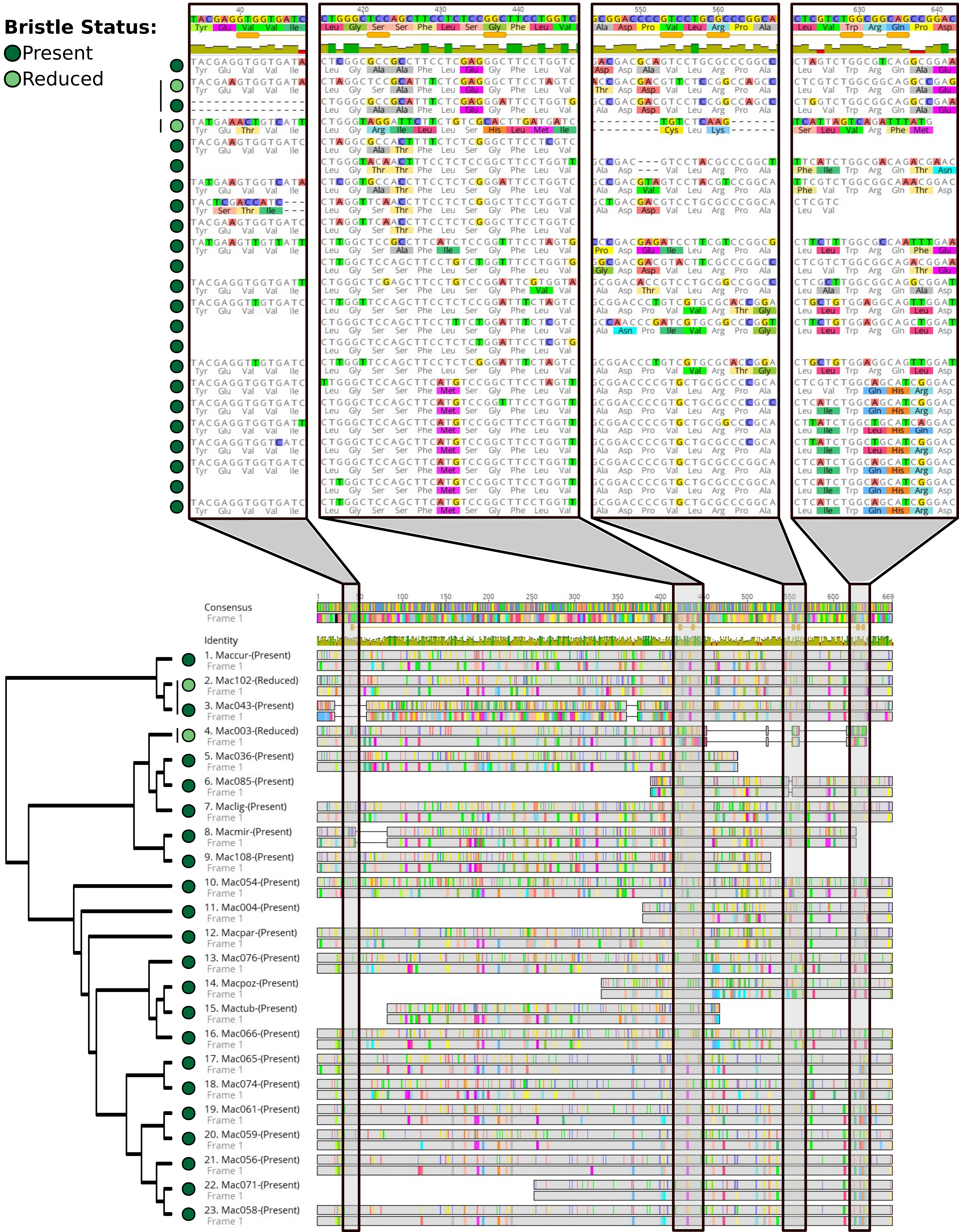
