## Supplementary figures and images for "Faster rates of molecular sequence evolution in reproduction-related genes and in species with hypodermic sperm morphologies"

### Figure S10

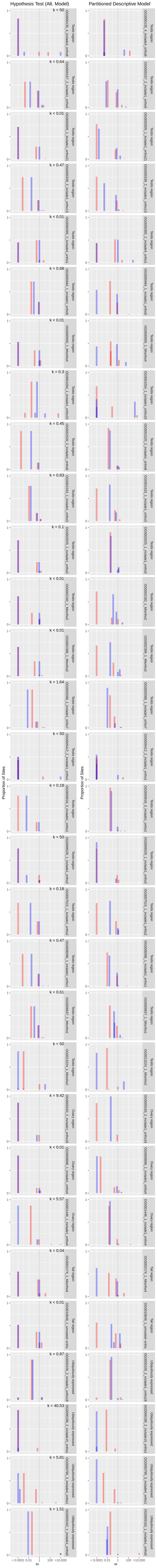

### Figure S11

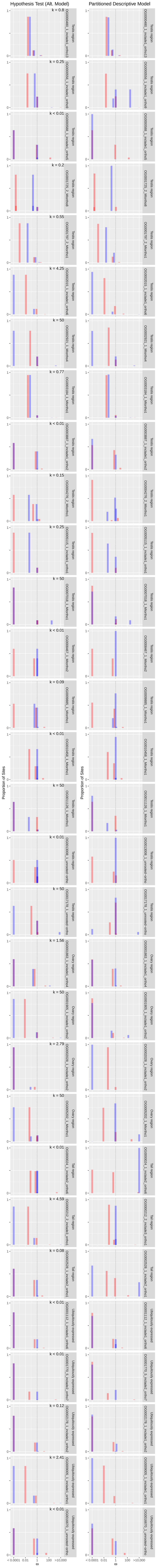
