## Supplementary Material for "Faster rates of molecular sequence evolution in reproduction-related genes and in species with hypodermic sperm morphologies"

Wiberg et al.

Bioinformatic notes of the analysis pipeline, along with shell and R scripts as well as Supplementary Archive 1, which includes all final alignments and pruned trees used in the analyses, can be found in a separate Zenodo repository: https://doi.org/10.5281/zenodo.4972188

**Supplementary Text**

**T1. Parametric statistics for results of three-ratio branch models**

In spite of the fact that ω values showed skewed distributions and unequal variances between the bristle states, we also wanted to explore the robustness of the analyses presented in the main text, by testing the two factors “bristle state” and “annotation group” simultaneously, including an interaction term, using two approaches. We $\log_{10}()$ transformed all ω values and fit a repeated measures MANOVA using the “MANOVA.RM” (v. 0.4.3; Friedrich et al., 2021) R package where the three ω values from the same OG are considered repeated measures since they are not independent. We found that there was an overall effect of the annotation group (Wald-Type $\chi^2$ = 21.6, d.f. = 3, $p < 0.001$) and of bristle state (Wald-Type $\chi^2$ = 54.8, d.f. = 2, $p < 0.001$) on ω values, but no effect of the interaction (Wald-Type $\chi^2$ = 6.7, d.f. = 6, p = 0.35). Similar results were obtained if we fit a linear mixed effects model with the “nlme” (v. 3.1-149; Pinheiro et al., 2020) R package, treating OG as a random effect to control for the non-independence of ω values within OGs. Again, there was an overall effect of the annotation group ($F_{3,510} = 6.4$, $p < 0.001$) and of bristle state ($F_{2,1020}$ = 24.3, $p < 0.001$) on ω values, but no effect of the interaction ($F_{2,1020} = 0.6$, p = 0.76). These results therefore recapitulated the trends we described in the main text using separate non-parametric statistical tests.

**T2. Removal of residual cDNA synthesis primer sequences**

We discovered in Brand et al., (*in press*) that the transcriptome assemblies used in this analysis contained residual cDNA synthesis primer sequences. For full details see Brand et al., (*in press*). Briefly, a 21nt sequence corresponding to a partial SMART-Seq v4 adapter could be found at the start of many assembled transcripts (between 11.7% – 70.9% across all assembled transcriptomes) derived from libraries produced using the SMART-Seq v4 Ultra Low Input RNA Kit. Despite the extensive alignment cleaning and filtering steps performed during the analysis steps of this manuscript, this sequence was also found in 47 alignments of CDS sequences that were analysed with PAML and RELAX (Testis region, 20; Ovary region, 5; Tail region, 3; and Ubiquitously expressed, 19, out of a total of 1,149). This corresponds to ~4% of all analysed CDS alignments. In order to test whether the inclusion of these alignments greatly influences the results, we repeated all analyses excluding these 47 alignments and found that results were not qualitatively different. We are therefore confident that this cDNA primer is not systematically influencing our estimates of ω and throughout the main text we present the results including all alignments.

**T3. Description of selected candidate orthogroups**

The tail-region annotated orthogroup (OG) “OG0003507_1_inclade1_ortho3” (from Mlig022958.g1), includes four independent losses and two independent reductions in a fairly good alignment. It shows $\omega$ higher than the median for ubiquitously-expressed OGs in species of all bristle states, and evidence for positive selection from branch-site tests in the present-reduced contrast. Despite its annotation, *in situ* hybridisation screens suggest that this transcript (called RNA815_50706 in an earlier transcriptome version) is mainly expressed in the testes, but also in the gut (Weber et al., 2018).

An example with more detailed functional information can be seen in the testis-region annotated “OG0000305_1_inclade1_ortho17” (from Mlig020950.g1). This gene has higher $\omega$ than the median for ubiquitously-expressed OGs, but only for species with reduced bristles, and there is no evidence for positive selection from branch-site tests for either the present-absent or present-reduced contrast. However, it shows evidence of relaxed selection in the RELAX analyses (present-absent contrast). Knockdown experiments of this gene (also called *Mlig-sperm1*) give an aberrant sperm phenotype and result in reduced fertility (Grudniewska et al., 2018). Thus, one explanation is that this gene is crucial in species with present bristles to maintain morphology, but is under relaxed selection (reduced intensity of selection) in species with absent or reduced bristles because it is no longer needed. This may also help to explain why there are very few representative sequences from species with absent bristles.

The gene “OG0000247_1_inclade3_ortho5” is also annotated as testis-region specific (from Mlig045906.g1). It shows a nice alignment and a higher $\omega$ than the median for ubiquitously-expressed OGs, but only for species with present or reduced bristles. Additionally, it shows evidence for positive selection from branch-site tests (present-absent contrast).

The tail-region annotated “OG0010649_1_unrooted-ortho” (from Mlig009359.g1; n = 7 independent losses; table S1 and S2) has higher $\omega$ than the median for ubiquitously-expressed OGs in all bristle states, and it is identified as having a few sites under positive selection in branch-site tests (though not after strict correction for multiple testing; table S1). The alignment is also well behaved, with long blocks of well-aligned codons. From *in situ* hybridisation screens in *M. lignano* by Weber et al. (2018) this transcript (RNA815_16384 in earlier transcriptome) is known to be expressed specifically in the stylet, which is one of the morphological characters that is strongly associated with the mating strategy and the sperm morphology.

Finally, the tail-region annotated “OG0002759_1_inclade1_ortho4” (from Mlig003431.g4; RNA815_23759 in earlier transcriptome) is expressed in the prostate glands, ovaries, developing eggs, and testes (Weber et al., 2018). This OG also has a good alignment and shows higher $\omega$ than the median for ubiquitously-expressed OGs in all bristle states. However, there is no evidence for

81 positive selection at specific sites from the branch-site test in either the present-absent or the present-reduced contrast.

### Supplementary Figures and Legends

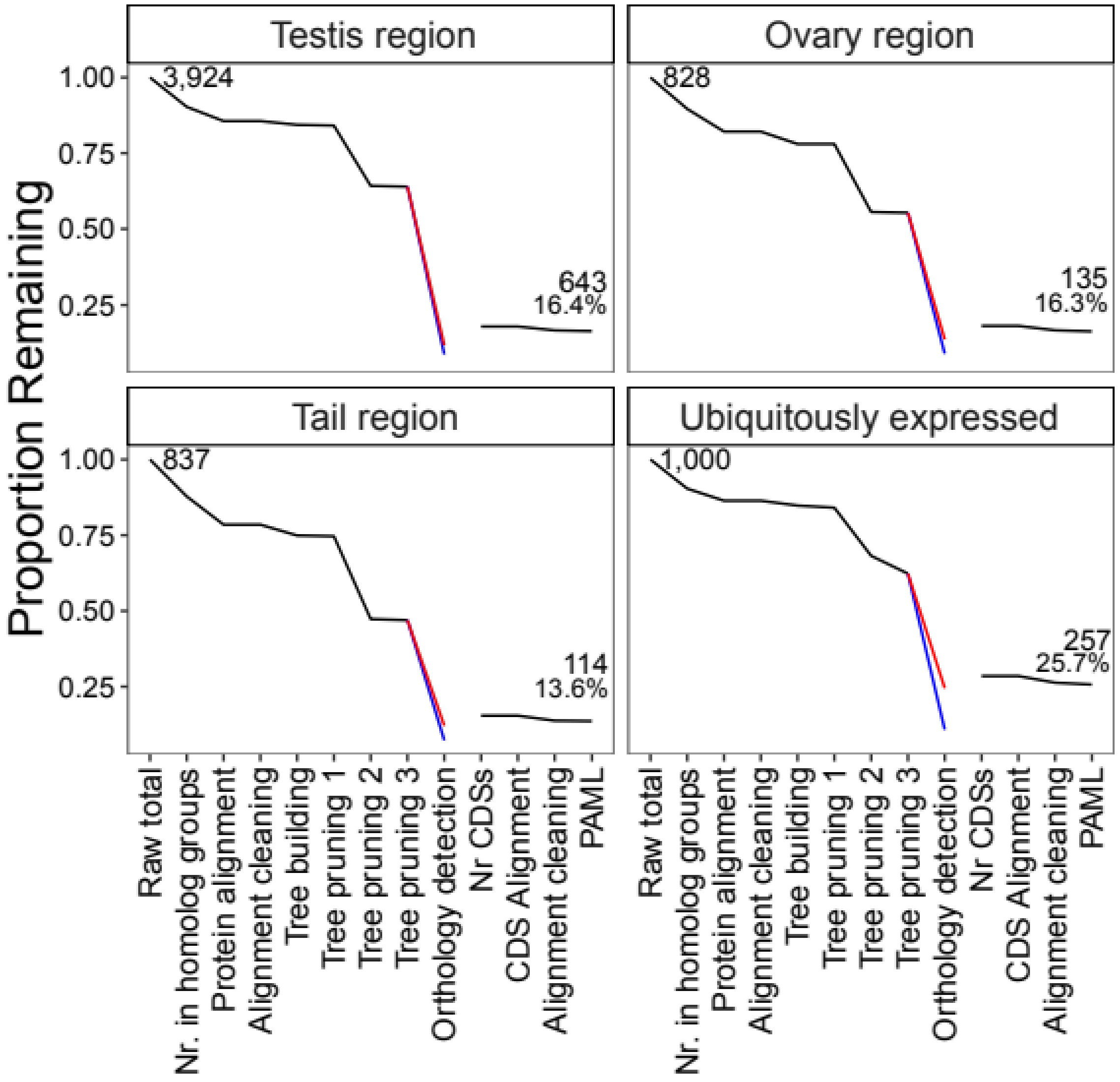


**Figure S1.** The proportion of *M. lignano* reproduction-related genes and ubiquitously-expressed genes (out of 1000 such genes initially selected at random) that remain after successive steps of the ortholog detection pipeline. Steps involving the ortholog inference algorithms are split into the “maximum inclusion” (MI; blue) and “rooted ingroups” (RT; red) algorithms. The gap between “Orthology detection” and “Nr CDSs” represents the fact that the final set of orthogroups (OGs) used is a union of the MI and RT OGs. The initial number of *M. lignano* genes for each group is given in the top right corner of each panel and the value in the lower left of each panel gives the final number (and percentage).

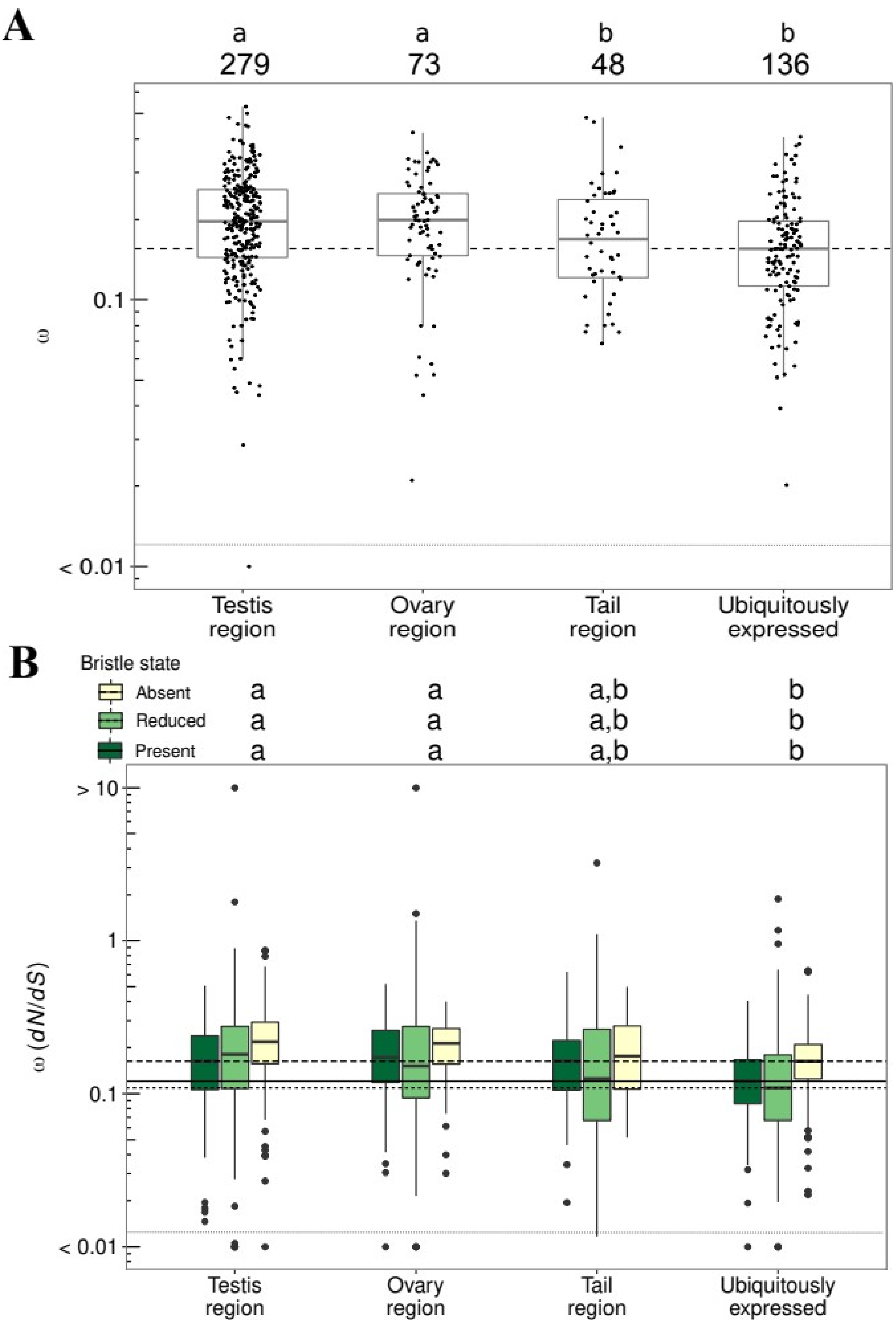


**Figure S2.** Analyses as in Figure 2, but data have been subset to exclude OGs that contain genes with < 3 independent losses of bristles. **A** *dN/dS* (ω) values, from a model assuming a single ω value for the whole tree, for orthogroups (OGs) assigned to different annotation groups. The horizontal dashed line gives the median ω value for the ubiquitously expressed group. Sample sizes for each annotation group are given as inset text. There was a significant effect of annotation group on average ω (Kruskal-Wallis rank-sum test, $\chi^2 = 27.8$, d.f. = 3, $p < 0.001$) **B** *dN/dS* (ω) across orthogroups (OGs) in different annotation groups for species with different sperm morphologies (bristle state). The solid, short-dashed, and long-dashed horizontal lines give the median ω values across OGs annotated as ubiquitously expressed for species with different sperm morphologies (see inset legend). For all bristle states there was a significant effect of annotation group on average ω (Kruskal-Wallis rank-sum tests, Present: $\chi^2 = 22.5$, d.f. = 3, $p < 0.001$, Reduced: $\chi^2 = 34.1$, d.f. = 3, $p < 0.001$, Absent: $\chi^2 = 32.4$, d.f. = 3, $p < 0.001$). For both **A** and **B** the letters show nonparametric all-pairs post-hoc tests (groups with different letters are significantly different at an adjusted p-value < 0.05), and for ease of plotting, some OGs with ω estimates > 10 or < 0.01 have been fixed to ω values of 10 or 0.01, respectively, and separated from the remaining points with stippled horizontal lines.

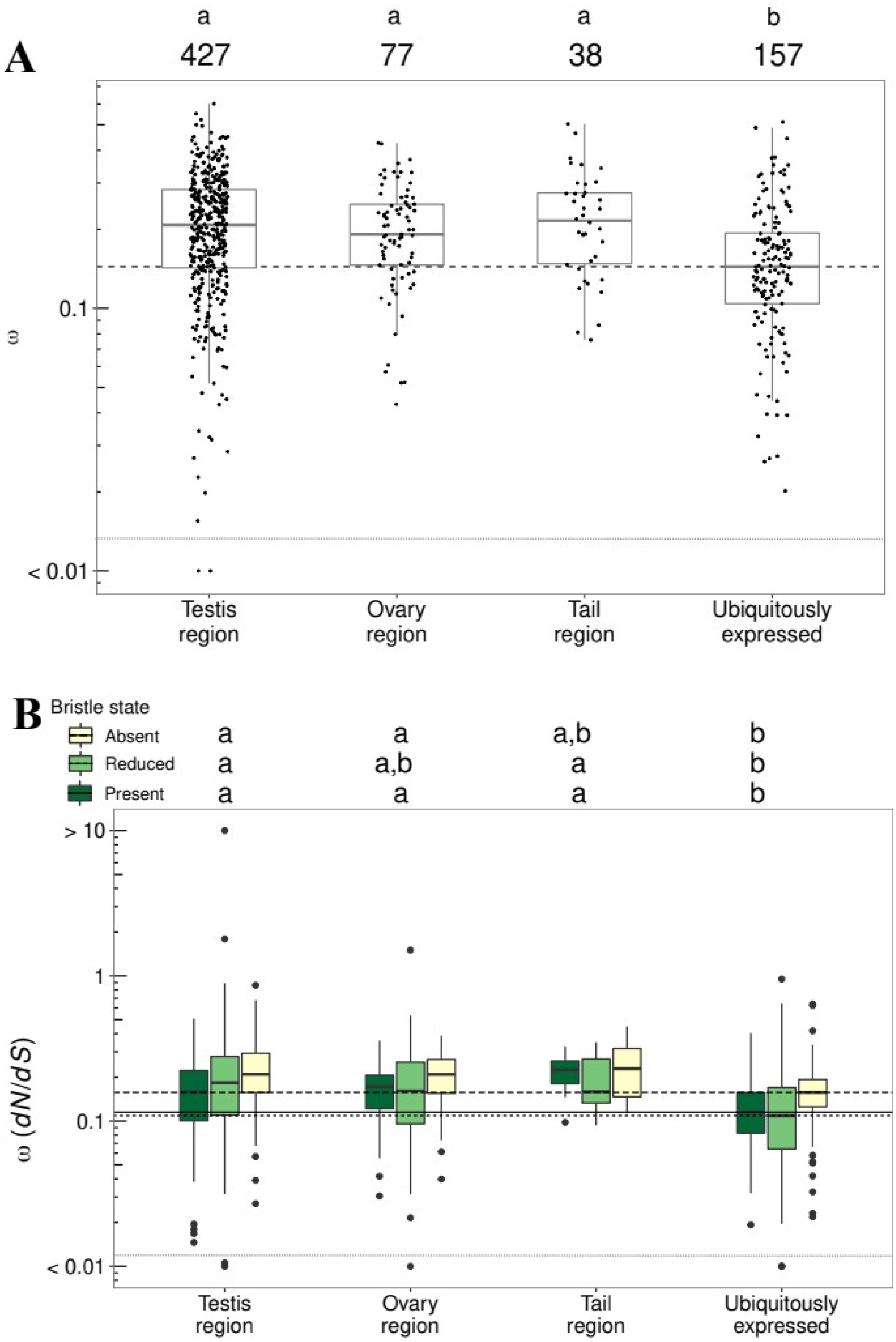


**Figure S3.** Analyses as in Figure 2, but data have been subset to exclude OGs that contain genes with low expression in *M. lignano* (< 50 mapped reads). **A** *dN/dS* (ω) values, from a model assuming a single ω value for the whole tree, for orthogroups (OGs) assigned to different annotation groups. The horizontal dashed line gives the median ω value for the ubiquitously expressed group. Sample sizes for each annotation group are given as inset text. There was a significant effect of annotation group on average ω (Kruskal-Wallis rank-sum test, $\chi2 = 42.7$, d.f. = 3, $p < 0.001$) **B** *dN/dS* (ω) across orthogroups (OGs) in different annotation groups for species with different sperm morphologies (bristle state). The solid, short-dashed, and long-dashed horizontal lines give the median ω values across OGs annotated as ubiquitously expressed for species with different sperm morphologies (see inset legend). For all bristle states there was a significant effect of annotation group on average $\omega$ (Kruskal-Wallis rank-sum tests, Present: $\chi^2 = 27.8$, d.f. = 3, $p < 0.001$, Reduced: $\chi^2 = 26.8$, d.f. = 3, $p < 0.001$, Absent: $\chi^2 = 30$, d.f. = 3, $p < 0.001$). The only difference compared to the analyses in Figure 2B was that for species with reduced bristles, there was now also a significant difference between OGs with tail-region annotations and OGs annotated as ubiquitously expressed. For both **A** and **B** the letters show nonparametric all-pairs post-hoc tests (groups with different letters are significantly different at an adjusted p-value < 0.05), and for ease of plotting, some OGs with $\omega$ estimates > 10 or < 0.01 have been fixed to $\omega$ values of 10 or 0.01, respectively, and separated from the remaining points with stippled horizontal lines.

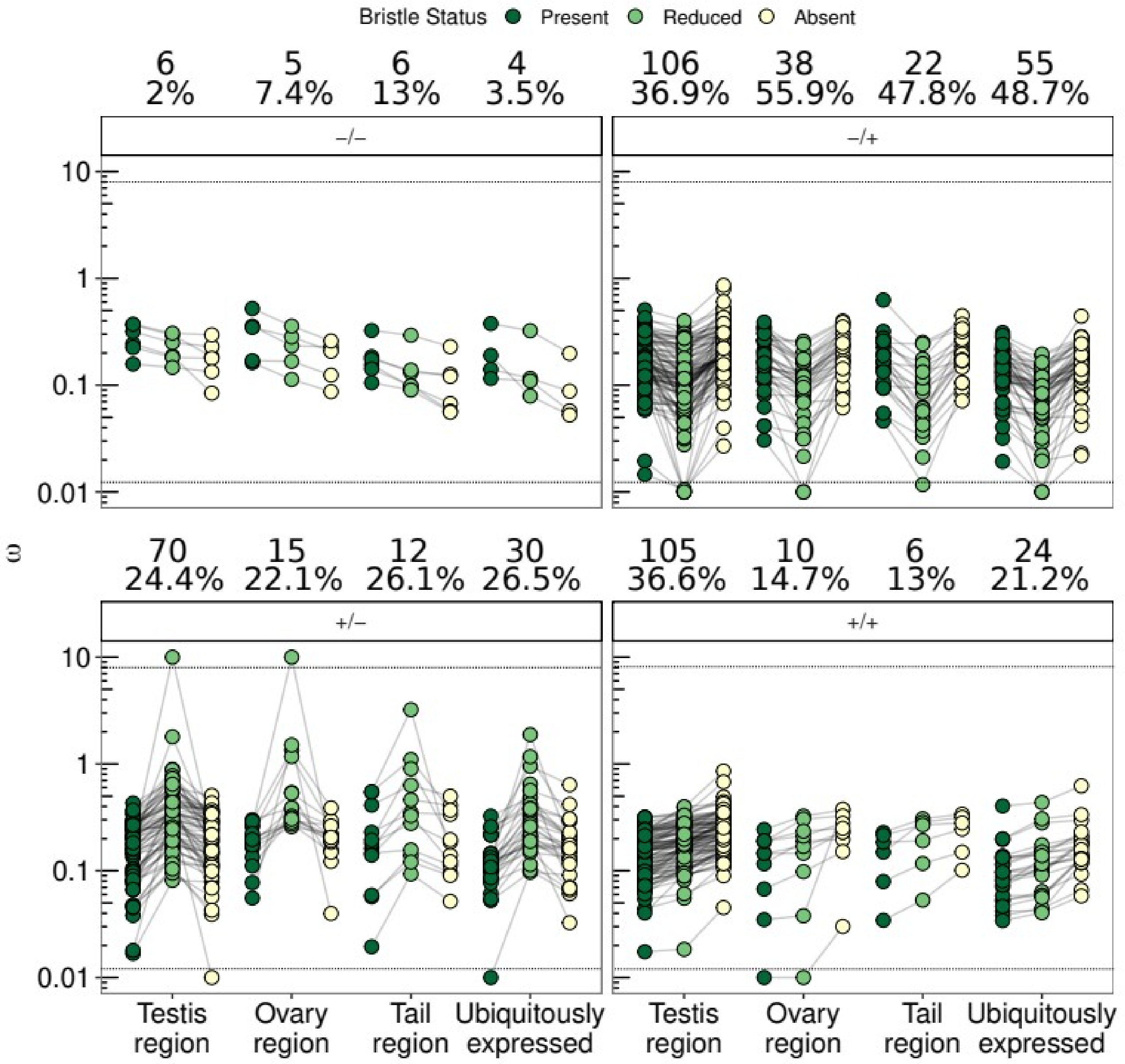


**Figure S4.** *dN/dS* (ω) values for orthogroups (OGs) that were better explained by a model assuming different ω values for species with the three contrasting sperm morphologies (bristle states). Values for the same OG in species with different bristle states are connected by a line. Panels show the data split into different patterns, namely where ω values always decrease (-/-) or increase (+/+), as well as two (-/+ and +/-) where species with reduced bristles do not show intermediate ω values, as one moves from species with bristles present, *via* those with reduced bristles, to those with absent bristles. The text above each panel gives the number and percentage of OGs across annotation groups and patterns of ω. Note that for ease of plotting, some OGs with ω estimates > 10 or < 0.01 have been fixed to ω values of 10 or 0.01, respectively, and separated from the remaining points with stippled horizontal lines.

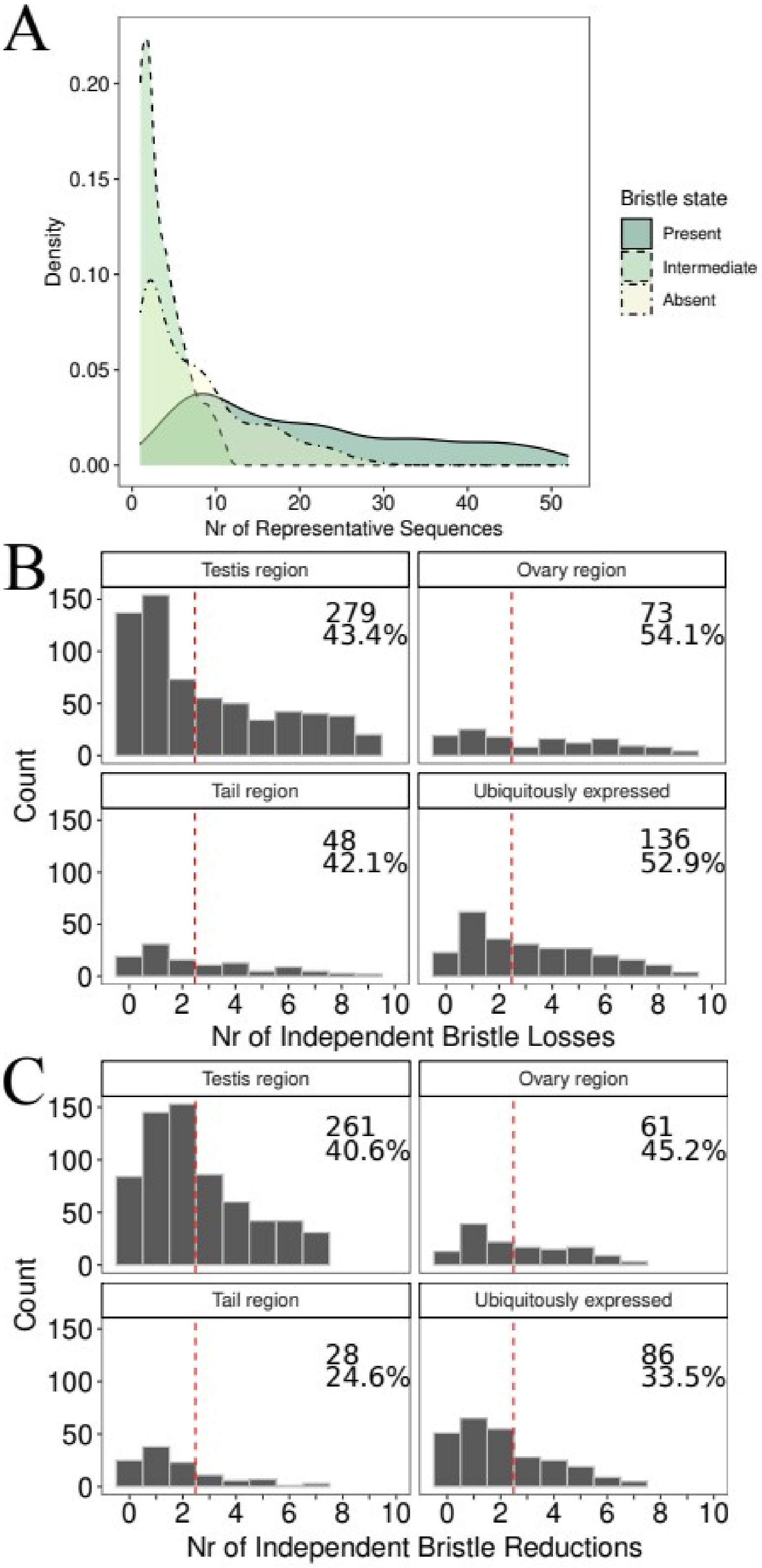


**Figure S5.** The distribution of the number of representatives from each bristle state across the orthogroups (OGs). **A** The number of representative sequences from species with different sperm bristle states across OGs. **B** The number of independent bristle losses across OGs in different annotation groups. **C** The number of independent bristle reductions across OGs in different annotation groups. The inset values in **B** and **C** give the number and percentage of OGs beyond the dashed red line (i.e. ≥3).

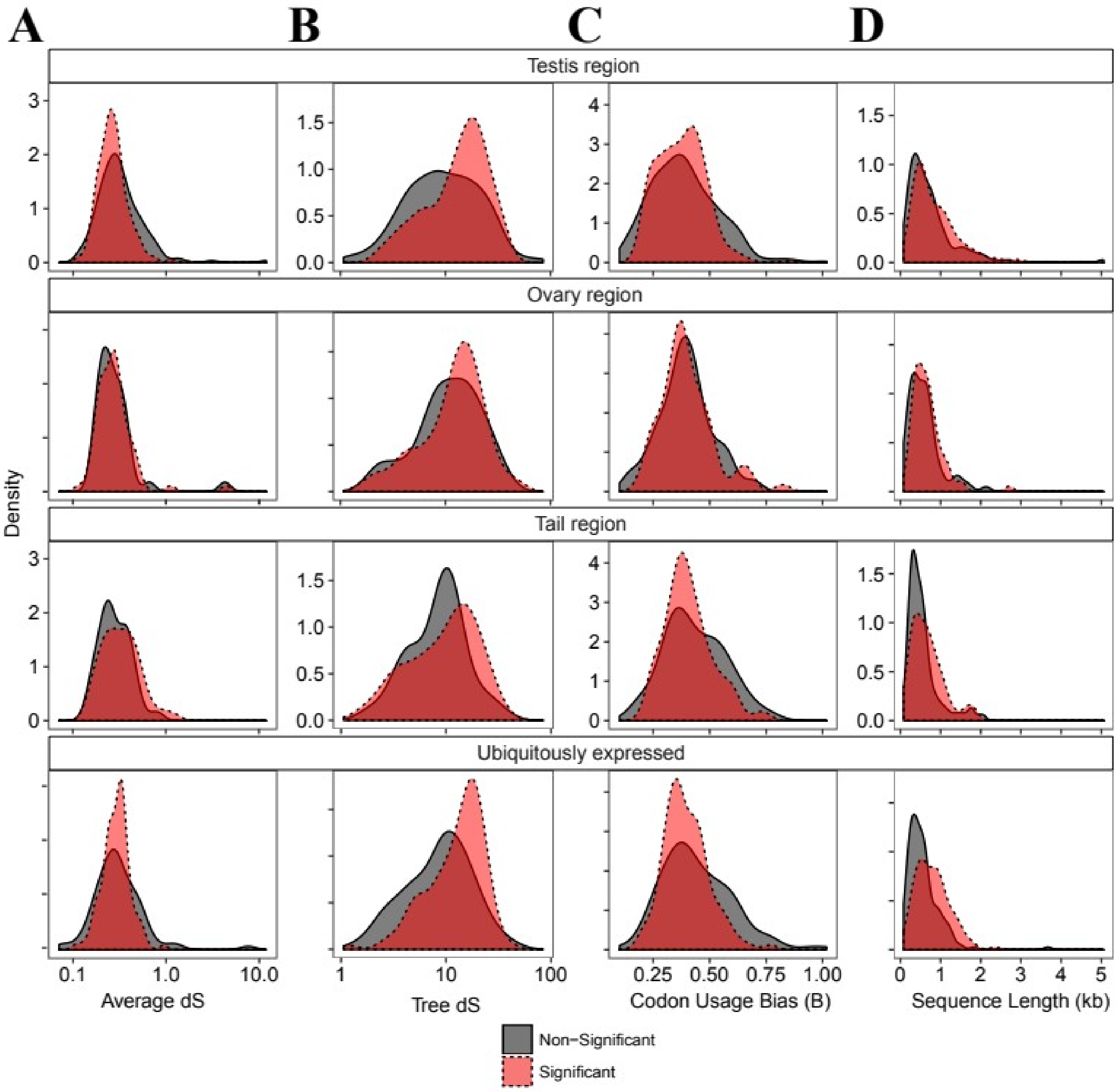


**Figure S6.** Comparisons between the orthogroups (OGs) that either significantly support a model with three separate ω values (red, significant) or are better represented by a model with a single ω value (grey, non-significant). **A** Average pairwise *dS* of terminal branches. **B** total tree *dS* (t*he summed dS* of all branches in the tree). **C** codon usage bias of species with absent bristles compared to species with bristles present. **D** total OG length (in kb). All distributions are given for OGs that are significant (red) and non-significant (grey) for the alternative three ω model in PAML branch tests.

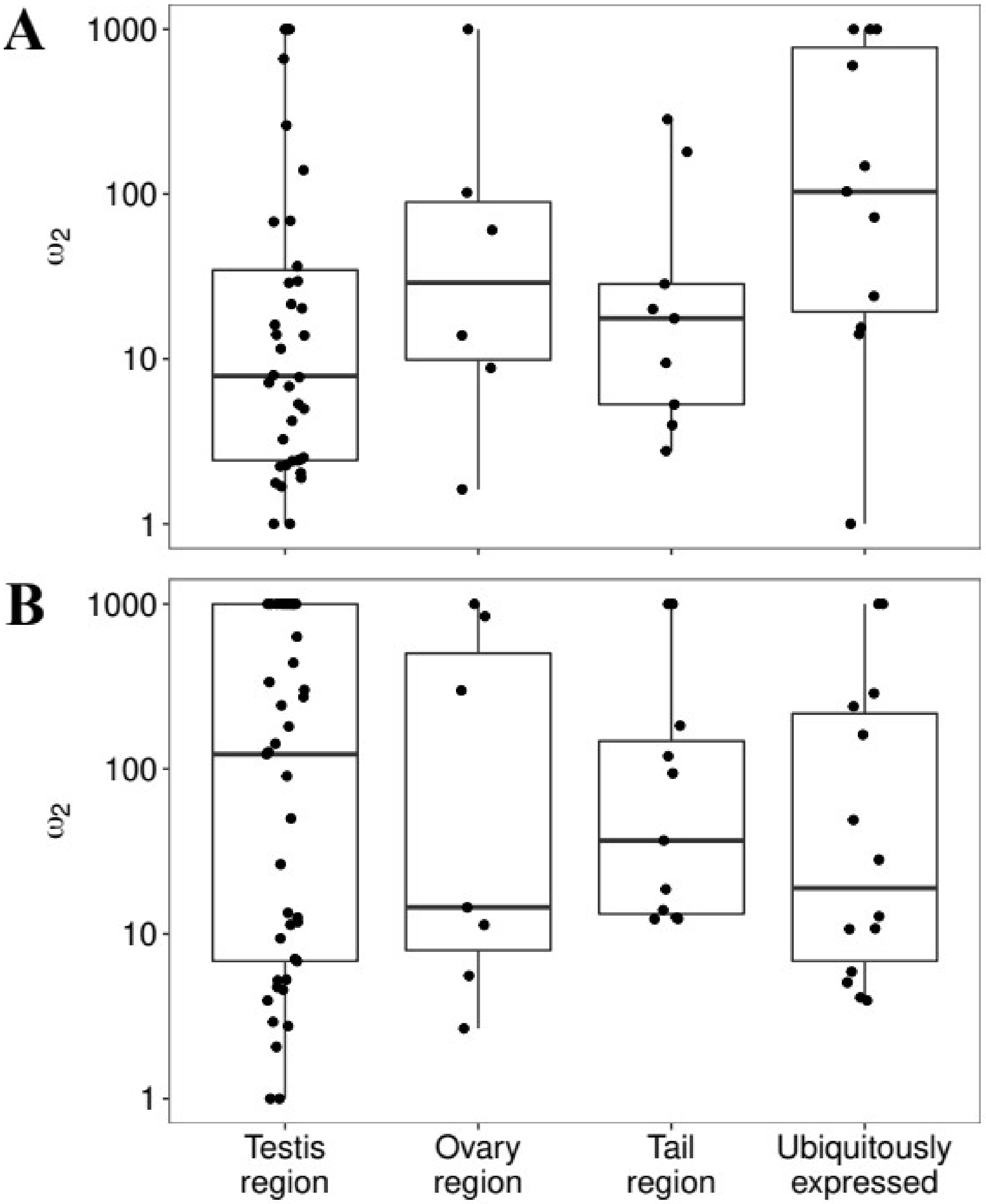


**Figure S7.** Boxplots of the value of the parameter $\omega_2$ in foreground branches from the branch-site model A for OGs in **A** the present-absent contrast and **B** the present-reduced contrast.

**Figure S8. (see separate .pdf file).** Alignments of selected orthogroups (OGs) with evidence for sites under positive selection contrasting species with bristles to species with absent or reduced bristles. Each page corresponds to a separate alignment. Sub-panels include the full alignment at the bottom of each page, as well as foci on regions containing sites determined to be under positive selection in PAML, with a yellow and red marker indicating a Bayes Empirical Bayes posterior probability, BEB $p > 0.95$ and $0.90 <$ BEB $p \leq 0.95$, respectively. The pruned phylogeny is presented next to the sub-panel of the full alignment and clades with independent losses/reductions of bristles are highlighted with a vertical line. The text at the top of each panel gives the label of the OG, the number of independent losses/reductions, the annotation group of the OG, as well as a legend for the bristle status.

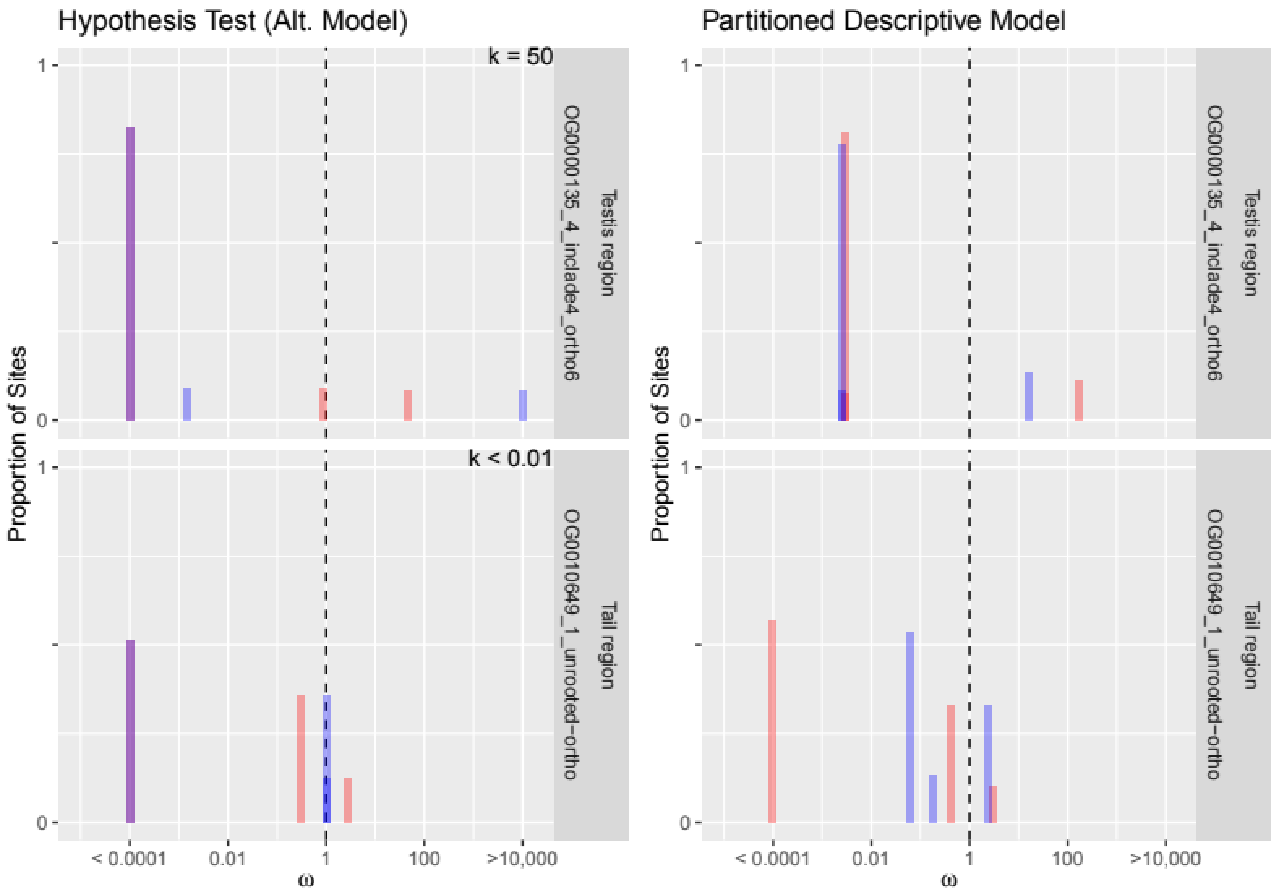


**Figure S9.** The proportion of sites within each site category that described the modelled distribution of ω (such that $\omega_0 \leq \omega_1 \leq 1 \leq \omega_2$) for reference branches (red bars, species with bristles) and test branches (blue bars, species with absent bristles) in the RELAX analyses. Here we show the OGs with highest and lowest estimated *k* values to aid the interpretation of the more detailed figures (see below). The left column shows the parameter estimates from the hypothesis testing framework comparing the null model to the alternative model. The right-hand column shows the parameter estimates from the less constrained Partitioned Descriptive Model (PDM; see Materials and Methods for details). For the top-left panel one can see that for test branches the distribution is wider than for the reference branches. Thus, the value of $k > 1$ indicates intensified selection. This pattern of a wider distribution for test branches is less clear, but can also be seen in the panel for the PDM estimates (top-right panel). The blue bars to the left of $\omega = 1$ have moved further to the left compared to the red bars. Meanwhile, in the bottom-left panel the distribution for test branches has become more narrowly distributed around 1 than the distribution for the reference branches. Thus, the value of $k < 1$ indicates relaxed selection in the test branches. From the PDM estimates there are clearly more sites with a value of $\omega \approx 1$ for test branches (i.e. a narrower distribution). For a larger set of examples, see figures S10 (present-absent contrast) and S12 (present-reduced contrast). Note that for ease of plotting, values < 0.0001 and > 10,000 have been set to 0.0001 and 10,000 respectively.

**Figure S10. (see separate .pdf file).** The proportion of sites within site categories that described the modelled distribution of $\omega$ (such that $\omega_0 \leq \omega_1 \leq 1 \leq \omega_2$) for reference branches (red bars; species with bristles present) and test branches (blue bars; species with absent bristles) in RELAX analyses. Shown for closer examination are the OGs with the 5 highest (rows 1-5) and 5 lowest (rows 6-10) values of *k,* as well as 20 random (rows 11-20) OGs. The left column gives the estimates from the alternative model in the null hypothesis testing framework of RELAX, while the right column gives estimates from a more relaxed Partitioned Descriptive Model (see Materials and Methods). Inset text in the left column shows the estimated *k* value. Note that for ease of plotting, values < 0.0001 and > 10,000 have been set to 0.0001 and 10,000 respectively.

**Figure S11 (see separate .pdf file).** The proportion of sites within each site category that described the modelled distribution of $\omega$ (such that $\omega_0 \leq \omega_1 \leq 1 \leq \omega_2$) for reference (red bars; species with bristles) and test (blue bars; species with reduced bristles) branches in RELAX analyses. Shown for closer examination are the OGs with the 5 highest (rows 1-5) and 5 lowest (rows 6-10) values of *k,* as well as 20 random (rows 11-20) OGs. The left column gives the estimates from the alternative model in the null hypothesis testing framework of RELAX, while the right column gives estimates from a more relaxed Partitioned Descriptive Model (see Materials and Methods). Inset text in the left column shows the estimated *k* value. Note that for ease of plotting, values < 0.0001 and > 10,000 have been set to 0.0001 and 10,000 respectively.

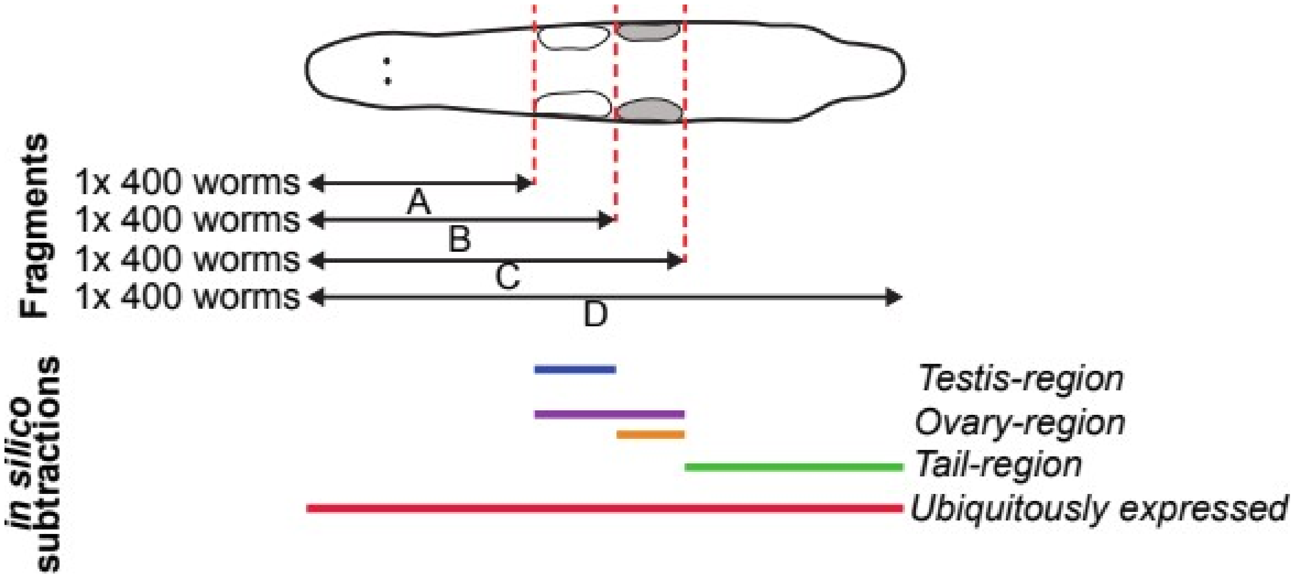


**Figure S12.** Schematic drawing of an adult worm indicating the cutting levels (red vertical dashed lines) used in Arbore et al. (2015) to produce four worm fragments (A, B, C, D). The expression patterns of each gene can be compared between neighbouring fragments to categorise them into reproduction-related annotation categories (testis-, ovary-, and tail-region). For example, a gene which increases (+) in expression between fragment A and fragment B, but then has no further increase in expression (0) between fragments B and C, as well as C and D, would obtain the pattern [+,0,0] and be labelled a "testis-region" gene. In a similar manner other patterns of expression define the remaining reproduction-related categories, namely ovary-region ([0,+,0] and [0,+,+]), and tail-region ([0,0,+]). Ubiquitously expressed genes are defined as those which did not differ in expression between any worm fragments (i.e. a pattern of [0,0,0], also called "non-specific" in Brand et al., 2020). For full details of these analyses see Arbore et al. (2015) and Brand et al. (2020). The figure is modified from Brand et al., (2020), worm drawing by J.N. Brand.
